# Haploinsufficient tumour suppressor PRP4K is negatively regulated during epithelial-to-mesenchymal transition

**DOI:** 10.1101/2020.04.19.043851

**Authors:** Livia E. Clarke, Allyson Cook, Sabateeshan Mathavarajah, Amit Bera, Jayme Salsman, Elias Habib, Carter Van Iderstine, Moamen Bydoun, Stephen M. Lewis, Graham Dellaire

**Author notes:** Corresponding Author: Graham Dellaire, Ph.D., Departments of Pathology and Biochemistry & Molecular Biology, Dalhousie University, P.O. BOX 15000, Halifax, Nova Scotia, Canada, B3H 4R2.

## Abstract

The pre-mRNA processing factor 4 kinase (PRP4K, also known as PRPF4B) is an essential gene. However, reduced PRP4K expression is associated with aggressive breast and ovarian cancer phenotypes including taxane therapy resistance, increased cell migration and invasion *in vitro* and cancer metastasis in mice; results consistent with PRP4K being a haploinsufficient tumour suppressor. Increased cell migration and invasion is associated with epithelial-to-mesenchymal transition (EMT), but how reduced PRP4K levels affect normal epithelial cell migration or EMT has not been studied. Depletion of PRP4K by small hairpin RNA (shRNA) in non-transformed mammary epithelial cell lines (MCF10A, HMLE) reduced or had no effect on 2D migration in the scratch assay but resulted in greater invasive potential in 3D transwell assays. Depletion of PRP4K in mesenchymal triple negative breast cancer cells (MDA-MB-231) resulted in both enhanced 2D migration and 3D invasion, with 3D invasion correlated with higher fibronectin levels in both MDA-MB-231 and MCF10A cells and without changes in E-cadherin. Induction of EMT in MCF10A cells, by treatment with WNT-5a and TGF-β1, or depletion of eukaryotic translation initiation factor 3e (eIF3e) by shRNA, resulted in significantly reduced PRP4K expression. Mechanistically, induction of EMT by WNT-5a/TGF-β1 reduced PRP4K transcript levels, whereas eIF3e depletion led to reduced PRP4K translation. Finally, reduced PRP4K levels after eIF3e depletion correlated with increased YAP activity and nuclear localization, both of which are reversed by overexpression of exogenous PRP4K. Thus, PRP4K is a haploinsufficient tumour suppressor negatively regulated by EMT, that when depleted in normal mammary cells can increase cell invasion without inducing full EMT.

## INTRODUCTION

Epithelial-to-mesenchymal transition (EMT) is an important process during embryonic development where cell-cell contacts are broken in concert with a loss of apical-basal polarity, and cytoskeletal reorganization, which converts immotile epithelial cells into motile mesenchymal cells (1). The process of EMT is regulated by transcription factors like Zeb1 that induce the expression of mesenchymal genes, and transcriptional factors Snail/SNAI1 and Slug/SNAI2 that repress the expression of epithelial genes such as E-cadherin (2). The dysregulation of EMT also plays an important role in cancer progression and metastasis; the latter being the leading cause of cancer-related death (1, 3). However, more recently partial or hybrid EMT states, where epithelial gene expression is maintained, have been described (4, 5). Partial EMT appears to have several advantages over complete EMT, including the ability to promote collective migration and invasion by cancer cells, resistance to cell death, and increased plasticity and metastatic potential, all of which contribute to increased tumorigenicity (4, 5). This is exemplified by the recent discovery that the maintenance of epithelial-like E-cadherin expression in mouse breast cancer models promotes cell survival during metastasis (6).

The overexpression of oncogenes and loss of genes encoding tumour suppressors are both linked to the incidence of cancer (7). Although the complete loss of a tumour suppressor gene is common in the development of cancer, incomplete loss, termed haploinsufficiency, can be sufficient for tumour development in some cases (8, 9). Several examples of haploinsufficient tumour suppressors exist, including the phosphatase PTEN (10), the ribosomal protein RPL5 (11), the transcriptional regulator CUX1 (12), the motor protein KIF1Bβ (13) and the acetyltransferase CREBBP (14). In these instances, there is typically 50% or greater loss of protein expression, yet even subtle changes in gene expression may affect tumour suppression. For example, just a 20% decrease in PTEN protein expression can promote the development of cancer (10).

In this study, we determined that the haploinsufficient tumour suppressor pre-mRNA processing factor 4 kinase (PRP4K, also known as *PRPF4B*) is negatively regulated by EMT. PRP4K is an essential kinase that is highly conserved across species (15, 16). PRP4K was initially described as *prp4* in *Schizosaccharomyces pombe*, which is a gene required for cell growth and regulation of pre-mRNA splicing in fission yeast (17, 18). Characterization of the PRP4K homolog in mammals determined that it associates with the U5 small nuclear ribonucleoprotein (snRNP), and is required for the formation of spliceosomal complex B during pre-mRNA splicing (16, 19). Although an essential gene in mammalian cells (20), reduced PRP4K expression is associated with poor outcomes in ovarian cancer (21). Furthermore, partial loss of PRP4K expression has been implicated in cellular processes that impact tumour suppression and chemotherapy sensitivity (15); these include regulation of the spindle assembly checkpoint (22), which impacts the cellular response to microtubule poisons including taxanes (23), transcriptional regulation (16, 24, 25), epidermal growth factor receptor (EGFR) signalling and anoikis (21), as well as Hippo/YAP signalling and cell migration (26). Therefore, PRP4K can be broadly classified as a haploinsufficient tumour suppressor. However, the impact of PRP4K depletion with respect to EMT-associated changes in gene expression, cell migration and invasion have not been investigated in normal epithelial cells.

Employing the non-transformed mammary epithelial cell lines MCF10A and HMLE, we determined that depletion of PRP4K by 50-90% using RNA interference leads to a partial EMT phenotype marked by increased 3D invasion (with generally no change or reduced 2D migration), reduced N-cadherin and no change in E-cadherin expression. In contrast, PRP4K depletion promoted both 2D cell migration and 3D invasion of the transformed triple-negative breast cancer cell line MDA-MB-231; with changes in 3D invasion in MDA-MB-231 and MCF10A cells correlating with increased fibronectin and Slug protein expression. In addition, we demonstrate that the induction of EMT in mammary epithelial cell lines promotes a decrease in PRP4K protein levels but the mechanism is dependent on the method of EMT induction. For example, induction of EMT by WNT5a/TGF-β1 treatment resulted in reduced PRP4K mRNA expression; whereas depletion of eIF3E, which is known to induce EMT in mammary epithelial cells (27), resulted in reduced PRP4K protein translation. Reduced PRP4K levels in eIF3E-depleted cells correlated with increased YAP activity and nuclear localization, which resulted in increased expression of YAP target genes (*ANKRD1, CTGF, CYR61*) and could be reversed by overexpression of exogenous PRP4K. Together, these data indicate that PRP4K is a haploinsufficient tumour suppressor negatively regulated by EMT, and that when depleted in normal mammary cells can increase cell invasion without inducing full EMT.

## MATERIALS AND METHODS

### Cell Culture

All cell lines used in this study were confirmed mycoplasma-free by an in-house testing regimen (MycoAlert, Lonza; LT07-703). MDA-MB-231 cells from the American Type Culture Collection (ATCC; Gaithersburg, MD, USA) were cultured in Dulbecco’s modified Eagle medium (DMEM; Sigma-Aldrich; Oakville, ON, CA) supplemented with 10% fetal calf serum, 1% penicillin/streptomycin at 37°C with 5% CO_2_. HMLE (gift of Dr. Shashi Gujar, Dalhousie University) and MCF10A (ATCC) cells were cultured in Dulbecco’s Modified Eagle’s Medium/Nutrient Mixture F-12 (Sigma-Aldrich), supplemented with reagents from Sigma-Aldrich including: 5% horse serum, 1% penicillin/streptomycin, 0.5 μg/mL hydrocortisone, 10 μg/mL insulin, 10 ng/mL epidermal growth factor at 37°C with 5% CO_2_. In addition, cholera toxin (1 ng/mL) (Sigma-Aldrich) was added to the MCF10A media.

For Actinomycin D treatment, cells were treated with 10 μg/ml Actinomycin D (Sigma-Aldrich) for the indicated time periods and then harvested for RT-qPCR analysis. For cycloheximide treatment, cells were treated with 50 μg/ml of cycloheximide (Sigma-Aldrich) for the indicated time periods and then harvested for Western blot analysis.

### Lentiviral Transduction

To generate the MDA-MB-231, HMLE and MCF10A TRIPZ shPRP4K cell lines TRIPZ (shPRP4K-1 = clone: V2THS_383962, shPRP4K-2 = clone: V3THS_383960, Non-silencing shCTRL = RHS4743) lentiviral shRNAs were purchased from ThermoFisher Scientific (Thermo; Mississauga, ON, CA). Lentivirus was produced by co-transfection of the TRIPZ shRNA with pMD2.G and psPAX2 (gifts from Didier Trono, Addgene plasmid #12259 and #12260, respectively) lentiviral packaging vectors into human HEK-293T cells via calcium-phosphate transfection (Promega; Madison, WI, USA), according to manufacturer’s directions. After 48h, media from the transfected cells was filtered using a 0.45μ filter, and the viral media added to the target cell line for 48h. Following the 48h, the cells were split and treated with virus for another 48h to increase the transfection efficiency. Cells were allowed to recover in fresh media for 24h followed by selection in fresh medium containing 2μg/ml puromycin for 4 to 5 days. To induce expression of the inducible PRP4K shRNA, 5μg/mL doxycycline (Sigma-Aldrich) was added to culture media for 96 h prior to experimentation with the drug being replaced every 24 h.

### Retroviral Transduction and Cell transfections

Production of MCF10A control and shEIF3E cell lines has been previously described (27). MCF10A shCtrl and shEIF3E cell lines were transfected in 100 mm tissue culture plates with 5 μg of DNA at a ratio of 1:50 of a T7-tagged PRP4K expression vector (16) or pBluescript SK-(Stratagene/Agilent Technologies, Santa Clara, CA, USA) using the Neon Transfection System (Thermo) according to the manufacturer’s directions. A pulse voltage of 1400V, pulse width of 20 ms and pulse number of 2 was used. Cells were harvested for RT-qPCR, fixed and permeabilized for immunofluorescence 36 h post-transfection.

### Induction of EMT by media supplementation

StemXVivo EMT Induction Media Supplement was purchased from R&D Systems (Oakville, ON, CA). Cells were plated in 10 cm plates (HMLE-5 x 10^5^ cells; MCF10A-1.5 x 10^6^ cells) with 6 ml of complete media. Sixty microliters of 100X StemXVivo EMT Induction Media Supplement was added to each plate. Three days following cell plating, media was removed and replaced with fresh media and supplement. Two days later, plates were harvested for Western blot and RT-qPCR analysis.

### Scratch Assay

Cells were seeded such that plates would be 90-100% confluent after 5 days. Twenty-four hours after plating the cells, doxycycline (5μg/mL; Sigma-Aldrich) was added to induce PRP4K knockdown and doxycycline was replaced every 24 hours. Seventy-two hours following doxycycline induction, media was replaced with low serum media (0.5%) to inhibit cell proliferation and 24 h later a scratch was made using a P200 pipette tip. Wounds made by the scratch were imaged every 4 h and overall imaging time before wound closure was dependent on the cell line, with data presented up to 48 h. Wound area was calculated using ImageJ software (NIH, Bethesda, MA, USA) and migration rate was determined as a percentage of wound area reduction (wound closure) over time.

### Transwell Assays

Growth factor-containing medium was added to the lower chamber and cells (MDA-MB-231: 5 x 10^4^ (Migration [M]), 1 x 10^5^ (Invasion [I]), HMLE: 1 x 10^5^ (M & I), MCF10A: 2 x 10^5^ (M & I)) reconstituted in serum-free medium were added to the upper chamber, which for migration assays were BioCoat™-Control-Cell-Culture inserts and for invasion assays were BioCoat™-Matrigel^®^ inserts (Corning, Bedford, MA, USA) of 24-well plates. After 24 hours, the non-migratory cells on the inside of the insert were removed. Inserts were then washed with PBS and fixed with methanol for 15 minutes. Following fixation, inserts were washed with PBS and stained with 0.5% crystal violet (Sigma-Aldrich) for 30 minutes. Inserts were then washed until clear of excess stain. Images of the inserts were taken using an Olympus CKX41 (Richmond Hill, ON, CA) equipped with ZEN lite 2012 acquisition software (Zeiss Canada, North York, ON, CA). For each replicate (n = 6), the number of migrating/invading cells in the insert was determined for 3-4 fields of view (2X magnification) and averaged.

### Western Blot Analysis

Cells were lysed in ice-cold lysis buffer (20mM Tris-HCl pH8, 300mM KCl, 10% Glycerol, 0.25% Nonidet P-40, 0.5mM EDTA, 0.5mM EGTA, 1x protease inhibitors) and lysates cleared by centrifugation (25min, 15 000xg, 4°C) before polyacrylamide gel electrophoresis and Western blotting as previous described^10^. Western blot densitometry was conducted using ImageJ software (NIH, Bethesda, MA, USA) and protein expression values were normalized against total protein or actin expression. Primary antibodies used for Western blot analysis include anti-beta-actin (Sigma-Aldrich, A2228) (1:10000), anti-beta-tubulin (Santa Cruz [Santa Cruz, CA, USA], sc-9104) (1:5000), anti-claudin-1 (Cell Signaling [Danvers, MA, USA], 13255) (1:1000), anti-E-cadherin (Cell Signaling, 3195) (1:10000), anti-eIF3e (Abcam [Toronto, ON, CA], ab32419) (1:1000), anti-Fibronectin (Abcam, ab32419) (1:1000), anti-N-cadherin (Santa Cruz, sc-59987) (1:1000), anti-PRP4K (Novus Biologicals [Oakville, ON, CA], NBP1-82999) (1:5000), anti-Slug (Cell Signaling, 9585) (1:1000), anti-Snail (Cell Signaling, 3879) (1:1000), anti-T7 (Millipore [Oakville, ON, CA], ab3790) (1:1000), anti-Trk-B (Santa Cruz, sc-377218) (1:1000), anti-Vimentin (Cell Signaling, 5741) (1:1000), anti-Yap1 (Sigma-Aldrich, WH0010413M1) (1:1000), anti-Zeb-1 (Cell Signaling, 3396) (1:1000), and anti-Zo-1 (Cell Signaling, 8193) (1:1000). Secondary antibodies used include HRP-conjugated sheep anti-mouse IgG (Sigma-Aldrich, A5906) and HRP-conjugated goat anti-rabbit IgG (Sigma-Aldrich, A6154).

### Immunofluorescence

Cells were plated onto sterile coverslips in 6-well plates and left to adhere overnight. Coverslips were washed with PBS for 5 minutes and then fixed in 4% paraformaldehyde for 30 minutes and permeabilized using 0.5% TritonX-100 (Sigma-Aldrich) for 5 minutes. Cells were washed again 3 times in PBS for 5 minutes each. Cells were blocked using 5% BSA (Bioshop; Burlington, ON, CA) in PBS for 20 minutes and then incubated with primary antibody for 1 hour at room temperature. Cells were washed 3 times with PBS for 5 minutes each and then incubated with Alexa Fluor secondary antibodies (Thermo) diluted 1:200 in 5% BSA (Bioshop) in PBS. Coverslips were washed 3 times with PBS for 5 minutes each, with 4’,6-diamidino-2-phenylindole (DAPI)(Sigma-Aldrich) added to the second wash at to a concentration of 300 nM to counterstain DNA. Coverslips were mounted on frosted glass microscope slides using Dako mounting medium (Dako North America, Inc; Carpinteria, CA, USA) and allowed to dry overnight in the dark at room temperature. Images of immuno-stained cells were acquired using a custom-built Zeiss Cell Observer spinning-disk laser confocal microscope (Intelligent Imaging Innovations, 3i; Boulder, CO, USA) equipped with a 63X 1.4 N.A objective (Zeiss) and a Prime95B scientific CMOS camera and Slidebook 6 software (3i). Primary antibodies for immunofluorescence include mouse anti-Yap1 (Sigma-Aldrich, WH0010413M1)(1:33) and rabbit anti-T7 (Millipore, ab3790)(1:100). Secondary anti-mouse Alexa Fluor 555 and anti-rabbit Alexa Fluor 647 antibodies (ThermoFisher) were used at a dilution of 1:200.

### RT-qPCR

Cell samples were lysed and homogenized using Trizol reagent (Thermo) according to the manufacturer’s directions and samples were frozen at −80°C for further analysis. RNA was isolated using the Ambion PureLink RNA Mini Kit (Thermo) according to the manufacturer’s protocol and included an on-column DNase I digestion. RNA quantity and quality were measured using a Nanodrop 2000 spectrophotometer (Thermo). Absorbance measurements A260/A280 and A260/A230 with ratios ~ 2.0 were accepted as pure for RNA. One microgram of RNA was reverse-transcribed to cDNA using the BioRad 5X iScript RT supermix kit (BioRad Laboratories Canada; Mississauga, ON, CA) for RT-qPCR, after which samples were diluted 1:1 with nuclease-free water. Samples without reverse transcriptase were included to confirm no genomic DNA contamination. Quantitative PCR (qPCR) was performed on cDNA samples using the 2X SsoAdvanced Universal SYBR Green Supermix (BioRad). A BioRad CFX Connect was used to perform the reactions and all experiments were done in triplicate. All primers used were designed using NCBI Primer Blast (https://www.ncbi.nlm.nih.gov/tools/primer-blast/). Gene expression data was normalized to at least two reference genes and analyzed using the BioRad CFX Maestro Software. Data were collected and analyzed as per the MIQE guidelines (28).

### Puromycin incorporation Assay

To measure relative rates of PRP4K translation we employed a puromycin incorporation assay that relies on the detection of protein puromycylation(29, 30). For this method, shCtrl and shEIF3E MCF10A cells were first grown in the absence of puromycin for 48 h prior to beginning the labelling with 10 μg/ml of puromycin for 2 h at 37 °C and 5% CO_2_. The cells were then harvested for detection of total puromycylated proteins as well as for immunoprecipitation (IP) using 20 μl (~2-3 μg) of sheep anti-PRP4K antibody H143 (16). IP was performed with the modified RIPA buffer. Briefly, after cold PBS washing, the MCF10A cells were lysed with IP lysis buffer (20 mM Tris, 350mM NaCl, 1mM EGTA, 1% NP-40) containing protease inhibitors for 30 min on ice. Next, lysed samples were centrifuged at 14 000 rpm for 20 min at 4 °C. After discarding the precipitate, the supernatants were collected and diluted to a final NaCl concentration of 150 mM to perform the IP. Cleared and diluted lysate was then incubated with the PRP4K antibody diluted in 0.5ml of PBS containing Dynabeads™ Protein G beads (Thermo) for 1 h, and incubated O/N at 4 °C. The next day precipitated complexes were washed with PBS two times, then washed with lysis buffer two times. The proteins were eluted with SDS sample buffer, boiled for 10 min and analyzed by SDS-PAGE as above, and nascent puromycylated proteins was detected by Western blot analysis as above using a mouse anti-puromycin antibody 12D10 (1:3000)(MABE343, Sigma-Aldrich).

### YAP Nucleo-Cytoplasmic Intensity Ratio Determination

Images were taken at random fields of view and then quantified. Two methods of quantification were done; a ratio of nuclear-cytoplasm YAP signal and the observed relative distribution of YAP localization in terms of increased nuclear signal (N>C), more cytosolic (N<C) and similar signals between nuclear and cytosolic (N=C). A minimum of 50 cells were quantified for each replicate (3 replicates total) to determine the ratios and relative distributions of YAP. Signals were measured using ImageJ and then ratios were determined or used to determine the relative distribution of YAP within the cell.

### Statistics

All graphs were made using GraphPad Prism 6. For conditions where two groups are being compared, a student’s t-test was used (two-tailed). For analysis of invasive breast carcinoma data in The Cancer Genome Atlas (31) (accessed *via* cbioportal.org on September 21, 2021), the logrank test was used to determine co-occurrence by comparing mRNA expression z-scores for selected genes, binned by high or low expression (>1 SD, or < −1 SD relative to diploid samples), and p-value estimates of significance were adjusted for false discovery rate to produce q-values (32). For experiments with three or more groups, a one-way ANOVA was carried out, employing Tukey’s post-hoc analysis to determine significance between groups. Analysis of scratch assay was carried out using linear regression to determine differences in the slope of wound closure percentage over time (i.e. rate of wound healing) between groups. Finally, for the analysis of nuclear (N) vs cytoplasmic (C) ratios (i.e. N>C, N=C, N<C) the Freeman-Halton extension of the Fisher exact probability test for a two-row by three-column contingency table was used employing an online tool: http://vassarstats.net/fisher2x3.html.

## RESULTS

### Depletion of PRP4Kpromotes invasion of both tumorigenic and non-transformed mammary cell lines but has differential effects on cell migration

Previously it had been shown that knockdown of PRP4K expression could increase the cell migration and invasiveness of the triple negative breast cancer (TNBC) cell line MDA-MB-231 (26); however, how PRP4K impacts cell migration and invasion of normal mammary epithelial cells has not been studied. Thus, we sought to determine whether loss of PRP4K had similar or differential effects on the migratory and invasive potential of breast cancer cells and non-transformed mammary epithelial cell lines. For these studies, we chose to deplete PRP4K by doxycycline-inducible short hairpin RNA (shRNA) in the normal mammary epithelial cell lines HMLE and MCF10A, and the triple negative breast cancer (TNBC) cell line MDA-MB-231; all of which have similar basal expression of PRP4K (Supplemental Figure S1). Across the 3 human breast cell lines, the level of PRP4K depletion ranged from 50 to 90%, with shPRP4K-2 having the greatest effect in each case. To assess the migratory potential of cells, we first employed the 2D scratch assay (Figure 1). Depletion of PRP4K increased the migratory potential of the MDA-MB-231 cell line in this assay as expected, but had no effect on the HMLE cells and significantly decreased migration in the MCF10A cells (Figure 1). These data indicate that the loss of PRP4K affects the migration of cells; however, the effect appears to be dependent on the transformation-state of the cell line, with more transformed and mesenchymal-like TNBC cells exhibiting increased 2D migration with loss of PRP4K.

**Figure 1.**
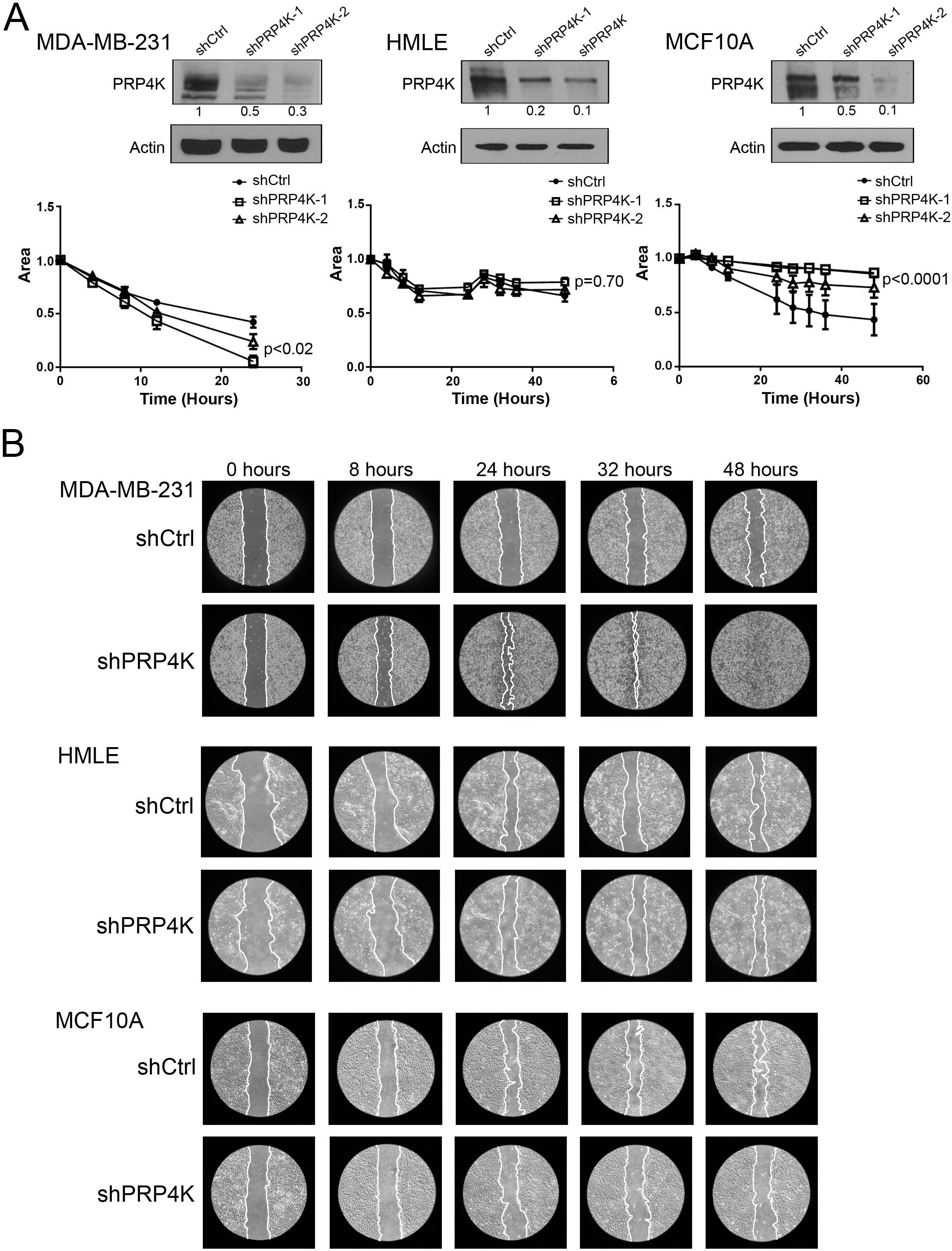
Effects of PRP4K depletion on the migration of normal mammary and transformed TNBC cells measured by wound healing scratch assay. (**A**) Normal mammary epithelial (MCF10A, HMLE) or TNBC MDA-MB-231 control (shCtrl) and PRP4K-depleted (shPRP4K-1 and shPRP4K-2) cells were grown to confluency and then a scratch was made using a P200 pipette tip and the average area of the scratch was set to a value of one and imaged every 4 h, and relative area of the scratched area plotted over 48 h. N = 3, error bars= SEM. Significance was determined by a linear regression comparing slopes between shCtrl and both shPRP4K hairpins. (**B**) Representative images of the scratched area measured in (A) shown over time.

We next conducted transwell assays with and without a matrigel-coated membrane to determine the invasive and migratory properties (respectively) of the cells following PRP4K depletion, respectively. In the absence of matrigel, the migratory potential of the MDA-MB-231 cell line in this assay was increased for one shRNA (shPRP4K-2) but decreased in presence of the other (shPRP4K-1) (Figure 2A). The migratory potential also decreased following PRP4K depletion in the HMLE cell line (Figure 3B), but increased with PRP4K reduction in the MCF10A cell line (Figure 2C). In the presence of matrigel, the invasive potential of the MDA-MB-231 (Figure 3A, shPRP4K-2) and MCF10A (Figure 3C; both shPRP4K-1 and −2) cell lines increased following PRP4K depletion. However, in the HMLE cell line, one shRNA with the greatest depletion of PRP4K (shPRP4K-2) significantly decreased the invasive potential of these cells (Figure 3B). Thus, depletion of PRP4K impacts 3D cell migration and invasion differently based on transformation-state and cell line. For example, depletion of PRP4K increases the migratory and invasive potential of MCF10A mammary epithelial cells in 3D culture, but decreases the migratory potential in 2D culture. This is in contrast to PRP4K-depleted MDA-MB-231 cells, which consistent with previous reports (26) showed increased migratory and invasive potential in both 2D and 3D culture.

**Figure 2.**
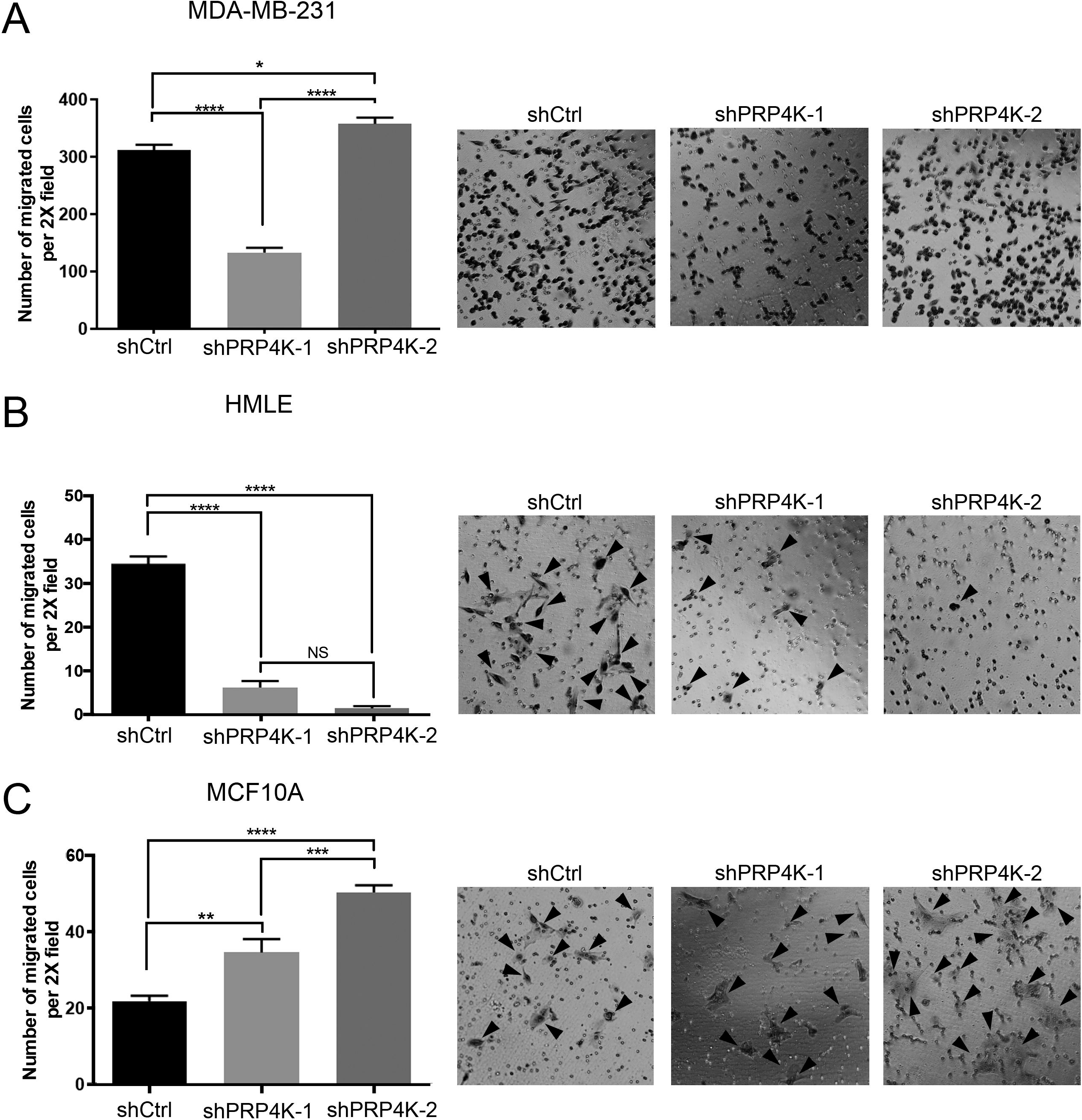
Effects of PRP4K depletion on the migration of normal mammary and transformed TNBC cells measured by transwell assay. (**A-C**) TNBC MDA-MB-231 or normal mammary epithelial (HMLE, MCF10A) control (shCtrl) and PRP4K-depleted (shPRP4K-1 and shPRP4K-2) cells were plated in transwells and migration into the insert membrane was determined 24 h later by visual enumeration and depicted as a bar graph (left), with representative images of each insert shown (right). Arrow heads indicate position HMLE and MCF10A cells that have migrated into the insert membrane. N = 6, error bars = SEM. Significance was determined by a one-way ANOVA. *p<0.05, **p<0.01, ***p<0.001, ****p<0.0001

### Depletion of PRP4K in non-transformed mammary epithelial cells is associated with altered epithelial and mesenchymal markers but does not induce full EMT

Recently we demonstrated that depletion of PRP4K in transformed mouse ID8 ovarian and human MCF7 breast cancer cells leads to dysregulation of EGFR signalling and upregulation of mesenchymal gene expression, including mesenchymal genes *Zeb1, vimentin* and *fibronectin* (21). From these data we hypothesized that reduced PRP4K expression might lead to cancer progression by promoting EMT and gene expression changes associated with EMT, which together may explain the differences in cell migration that we observed among the 3 mammary cell lines. Upon induction of PRP4K shRNA we observed no changes in E-cadherin expression in all three cell lines with both shRNAs, and a significant (or trending) increase in expression of mesenchymal markers slug (2 to 5 fold) and fibronectin (2 to 3 fold) in two out of three cell lines (MDA-MB-231 and MCF10A) with one or both shRNAs. We also observed cell and shRNA-specific changes in other mesenchymal markers including Snail (up in MDA-MB-231, down in HMLE and MCF10A), Zeb1 (up in MDA-MB-231 and HMLE), N-cadherin (down in HMLE and MCF10A) and vimentin (up in HMLE, down in MCF10A and MDA-MB-231) (Figure 4, Supplemental Figure S2). Similar variability between cell lines and shRNAs was seen for the epithelial markers Claudin-1 (up in MCF10A, down in MDA-MB-231 and HMLE), and Zo-1 (up in MDA-MB-231, and down in HMLE and MCF10A) (Supplemental Figure S2). Lastly, the mRNA expression of the same epithelial and mesenchymal factors did not consistently correlate with protein expression, suggesting post-transcriptional or post-translational control of these proteins following depletion of PRP4K (Supplemental Figure S3).

Overall, various mesenchymal markers were upregulated following PRP4K depletion in the 3 cell lines; with increased expression of fibronectin and Slug being the most reproducible gene expression change in PRP4K-depleted TNBC cell line MDA-MB-231 and non-transformed MCF10A cells. Analysis of TCGA data for invasive breast carcinoma (31), also revealed that low PRP4K mRNA expression is significantly correlated with high expression of fibronectin and slug in 526 breast cancers (Fig 4C). The expression of epithelial markers was also variable but importantly E-cadherin did not significantly change in response to PRP4K depletion in the 3 cell lines, with the overall effect being a mix of both epithelial and mesenchymal gene expression consistent with a “partial” EMT phenotype (5), particularly in the non-transformed mammary epithelial cell lines.

### EMT-induction media negatively regulates PRP4K gene expression

Given our data that depletion of PRP4K induces a partial EMT phenotype, which affects the 3D migratory and invasive potential of TNBC and non-transformed mammary epithelial cell lines, we next sought to determine whether the induction of EMT in non-transformed mammary epithelial cell lines affects the expression of PRP4K. Cells were cultured with an EMT-inducing media supplement consisting of various factors known to induce EMT (i.e. Wnt5a, TGFβ1, and antibodies against E-cadherin, sFRP1, and Dkk1)(33) in both HMLE and MCF10A mammary epithelial cells. Following treatment with the supplement, both HMLE and MCF10A cells became more spindle-shaped (Figure 5A), consistent with a more mesenchymal phenotype, and exhibited increased mesenchymal gene expression (Figure 5C). Furthermore, the protein expression of PRP4K was decreased in both cell lines as compared to untreated cells, and this correlated with decreased E-cadherin and increased N-cadherin protein and gene expression, consistent with classical or “complete” EMT (Figure 5B and 5C). In addition, we observed significant increases in the expression of other mesenchymal genes, including *fibronectin* and *vimentin* in both cell lines (Figure 5C).

To determine the mechanism of regulation for reduced PRP4K expression in response to EMT media we analysed both mRNA expression by RT-qPCR (Fig 6A) and protein-translation by measuring the incorporation of puromycin into nascent proteins (Fig 6B). This puromycin-incorporation assay has been used previously for assessment of translation by Western blot against puromycylated proteins (29, 30). The RT-qPCR analysis indicated that PRP4K mRNA expression was significantly reduced after 5 days treatment with EMT-induction media (Fig 6A). Although we could readily detect puromycylated proteins in control and EMT-media treated MCF10A cells, after immunoprecipitation of PRP4K there was no difference in translation when the ratio of puryomycylated PRP4K to total immunoprecipitated PRP4K was assessed (Fig 6B). Thus, these data indicate that complete EMT, marked by loss of E-cadherin and upregulation of N-cadherin, can be triggered by EMT-induction media (containing Wnt5a and TGF-β1) and results in decreased PRP4K mRNA expression.

### The induction of EMT by eIF3e depletion negatively regulates the translation of PRP4K

We also sought to induce EMT through another method that did not rely on a cocktail of supplements, which in turn would allow more extensive characterization of mechanisms underlying the negative regulation of PRP4K during EMT. To this end, we chose to induce EMT by depletion of eukaryotic translation initiation factor 3e (eIF3e) in MCF10A cells. EIF3e is a non-essential subunit of the eIF3 translation initiation complex (34, 35), and previously we demonstrated that depletion of eIF3e induces a robust EMT phenotype in the MCF10A cell line (27). When grown in culture, the MCF10A cells expressing a control shRNA (shCtrl) grew as tight clusters of well polarized epithelial cells (Figure 7A). Conversely, shRNA depletion of eIF3e using two different shRNAs induced a more mesenchymal phenotype characterized by single spindle-shaped cells, as previously described (27)(Fig 7A). Transcript and protein analysis of various epithelial and mesenchymal markers again confirmed that eIF3e depletion robustly induced EMT (Figure 7B and 7D). This EMT phenotype also correlated with a 50-60% decrease in PRP4K protein expression (Figure 7B), but in contrast to EMT-induction media did not significantly alter PRP4K (*PRPF4B*) mRNA expression (Fig 7C). Altogether, these data indicate that the induction of EMT by eIF3e depletion negatively affects PRP4K protein expression and that regulation of PRP4K occurs by either a post-transcriptional mechanism or at the level of protein translation.

Since the induction of EMT by depletion of eIF3e negatively affected PRP4K protein expression but had no effect on PRP4K transcript expression, we next sought to determine if PRP4K was being regulated in a post-transcriptional or post-translational manner. First, we examined the transcript stability of the *PRPF4B* gene encoding PRP4K in control and eIF3e depleted MCF10A cells treated with actinomycin D to inhibit RNA synthesis followed by RT-qPCR to assess *PRPF4B* transcript decay over time, and found no differences in turn-over of the *PRPF4B* mRNA (Figure 8A). Next, we examined the protein stability and turn-over of the PRP4K protein after treatment of control and eIF3e-depleted MCF10A cells with cycloheximide to block new protein translation, assessing protein levels over various time points (Figure 8B). Rather than seeing reduced levels of PRP4K overtime, the PRP4K protein appears to be stabilized after depletion of eIF3e compared to control MCF10A cells over the first 6 hours post-cycloheximide treatment (Fig 8B). These data indicate that during EMT induction by eIF3e depletion, PRP4K expression is not regulated at the level of mRNA or protein turn-over.

To assess how translation of PRP4K protein is altered during EMT induced by eIF3e depletion, we again employed the puromycin incorporation assay (Fig 8C). Consistent with previous studies (27), eIF3e-depleted cells exhibited reduced protein translation, as measured by reduced incorporation of puromycin in total protein compared to controls cells (Fig 8C). Upon immunoprecipitation, we detected a ~40% reduction in the puromycylation of PRP4K in the eIF3e-depleted cells compared to control when normalized to total immunoprecipitated PRP4K (Figure 8C). Thus these data indicate that, in the context of EMT induction by eIF3e depletion, changes in PRP4K protein expression are due to reduced translation.

### Depletion of eIF3e in MCF10A cells correlates with nuclear accumulation of YAP and activation of YAP-target genes, which is reversed by overexpression of PRP4K

Following the observation that the induction of EMT negatively affects PRP4K protein expression, we next sought to investigate the cellular consequences of PRP4K loss. Cho and colleagues recently demonstrated that PRP4K can negatively regulate YAP nuclear accumulation; an event that correlated with inhibition of YAP-target genes, many of which are involved in the development of cancer (26). Therefore, we hypothesized that negative regulation of PRP4K protein expression during EMT triggered by eIF3e depletion may similarly correlate with YAP nuclear accumulation and activation of YAP-mediated gene expression. To test this hypothesis, we examined the localization of YAP in control (shCtrl) and eIF3e-depleted MCF10A cells, with and without overexpression of exogenous PRP4K tagged with a T7 epitope (Figure 9). We found that the nuclear to cytoplasmic ratio of YAP was higher in eIF3e-depleted cells (Figure 9) and this correlated with increased expression of YAP target genes, including *CTGF, ANKRD1*, and *CYR61* (Figure 10A). This nuclear accumulation of YAP was dependent on PRP4K expression, as add-back of T7-PRP4K resulted in a significant reduction in the nuclear to cytoplasmic ratio of YAP (Figure 9). Overexpression of T7-PRP4K also reversed gene expression changes in YAP-target genes (i.e. *CTGF, ANKRD1* and *CYR61*) elicited by eIF3e-depleted MCF10A cells (Figure 10B). Thus, consistent with previous studies indicating that PRP4K regulates YAP nuclear accumulation and activity (26), loss of PRP4K during EMT correlates with an increased nuclear to cytoplasmic YAP ratio, and overexpression of PRP4K reverses this phenotype as well as changes in YAP-target gene expression in eIF3e-depleted MCF10A cells.

## DISCUSSION

For more than three decades it has been recognized that epithelial-to-mesenchymal transition (EMT) plays a key role in the acquisition of aggressive cancer traits that drive malignancy, such as the ability to invade tissue and metastasize. However, recent studies have linked aggressive cancer phenotypes to both mesenchymal-to-epithelial transition (MET) and developmental states caught “in-between” EMT and MET, otherwise known as partial EMT (1, 5, 36). In this study, we examined the inter-relationship between EMT and PRP4K, a protein that when depleted can induce aggressive cancer phenotypes including anoikis and chemotherapy resistance, as well as enhanced invasion potential and metastasis (21, 23, 26). Since the acquisition of complete or partial EMT often provides cells with increased migratory and invasive potential (5), we hypothesized that loss of PRP4K might also increase the migratory and invasive potential of both normal and breast cancer cells. Using the scratch assay to assess the 2D migratory potential of cells, we found that loss of PRP4K significantly increased the 2D migration of MDA-MB-231 TNBC cells (Figure 1). However, these results were in stark contrast to the migratory behaviour of the HMLE and MCF10A cell lines after PRP4K loss, where depletion of PRP4K significantly inhibited 2D migration of MCF10A cells and had little effect on the migratory behaviour of HMLE cells (Figure 1). Thus, the effects of PRP4K depletion on cell migration under 2D culture conditions appears to be cell line and transformation-state dependent, with little effect on normal epithelial cells and transformed and mesenchymal-like cells exhibiting increased 2D migration. We next examined the invasive and migratory potential of our cell lines in 3D culture conditions by employing transwell assays (Figure 2 and 3). In these studies, we found that depletion of PRP4K increased the migratory (Figure 2) and invasive potential (Figure 3) of both normal MCF10A mammary epithelial cells and TNBC cell line MDA-MBA-231 in 3D culture. However, both migration and invasion were inhibited in 3D culture conditions when PRP4K was depleted in HMLE cells. These data are not unique in showing a decoupling of migration in 2D and invasion properties in 3D culture conditions in relation to EMT, as previously it has been shown that induction of EMT in HMLE cells by overexpression of TWIST1 resulted in reduced 2D migration but increased invasion in the matrigel transwell assay (37).

**Figure 3.**
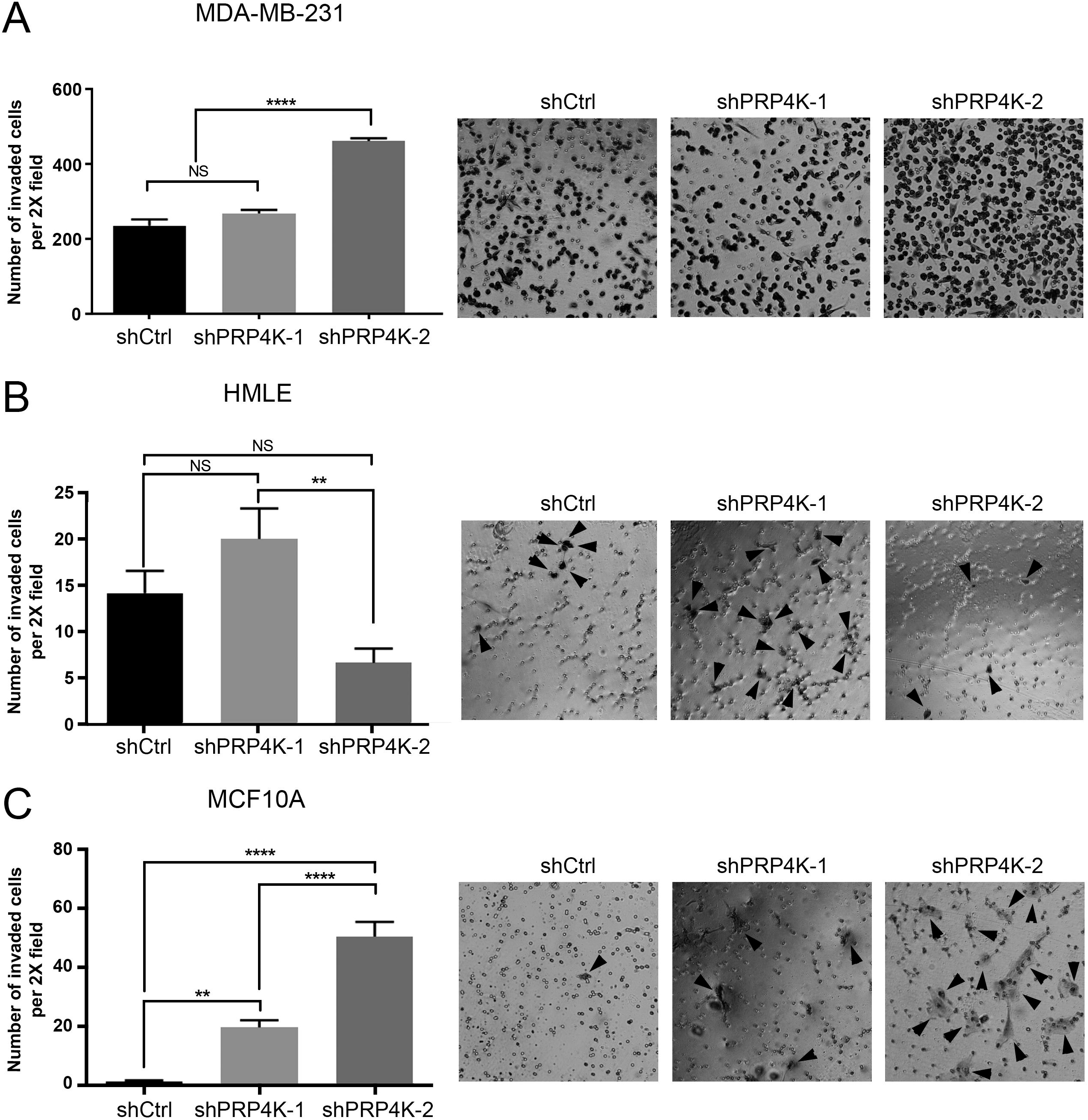
Effects of PRP4K depletion on the 3D invasion of normal mammary and transformed TNBC cells measured by matrigel transwell assay. (**A-C**) TNBC MDA-MB-231 or normal mammary epithelial (HMLE, MCF10A) control (shCtrl) and PRP4K depleted (shPRP4K-1 and shPRP4K-2) cells were plated in matrigel coated transwells and 3D invasion through the matrigel into the insert membrane was determined 24 h later by visual enumeration and depicted as a bar graph (left), with representative images of each insert shown (right). Arrow heads indicate position of HMLE and MCF10A cells that invaded the insert membrane. N = 6 experiments, error bars = SEM. Significance was determined by a one-way ANOVA. **p<0.01, ****p<0.0001.

Although our current study is the first to examine the impact of PRP4K depletion on the 2D and 3D migration and invasion behaviour of normal mammary epithelial cells, at least two other studies have looked at the effects of PRP4K depletion on cell migration using the MDA-MB-231 cell line. The first study, Cho *et al*., determined that siRNA mediated depletion of PRP4K increased migration in 2D employing the scratch assay over a 72 h time period, as well as increased invasion in 3D culture (26). In contrast, Koedoot *et al*., demonstrated reduced wound closure in the same 2D scratch assay over a 20 h time period when PRP4K expression was reduced by siRNA in MDA-MB-231 cells (38). Thus, our study is more aligned with the results of Cho and colleagues (26). Differences between these studies include the width of the “wound” generated during the 2D scratch assay, as well as the length of evaluation time (72 h vs 20 h). One interpretation is that with a wider wound the effects of PRP4K depletion on 2D migration may take at least 48 h to become apparent, as we observed complete wound closure at 48 h (Figure 3). A second possibility is the off-target effects and/or efficiency of PRP4K depletion by shRNA or siRNA in these experiments. Indeed, we observed more profound effects on migration and invasion correlating with greater reduction in PRP4K protein levels (i.e. shPRP4K-2 shRNA, Figure 2 and 3). Overall, these findings indicate that loss of PRP4K can increase the migration and invasion of an already-transformed cell line in both 2D and 3D culture, but only increases the migration and invasion of normal epithelial cell lines like MCF10A in 3D culture conditions.

Our data also indicate that the loss of PRP4K in “normal-like” mammary epithelial cell lines, such as MCF10A and HMLE, promotes a partial EMT phenotype instead of a complete EMT process. This was demonstrated by the upregulation of key mesenchymal proteins such as fibronectin and slug in MCF10A cells, while retaining expression of the key epithelial protein marker E-cadherin (Figure 4). Similarly, when PRP4K is depleted in the mesenchymal-like TNBC cell line MDA-MB-231, fibronectin and slug were upregulated but E-cadherin levels did not significantly change (Figure 4). Analysis of TCGA data for 526 invasive breast carcinomas (31), also indicated that low PRP4K expression in breast cancer is significantly correlated with high expression of fibronectin and slug (Fig 4C). In a previous study, exogenous addition of fibronectin to cultures of MCF10A cells was shown to promote EMT and increase migration and invasion on matrigel (39), which appears consistent with increased fibronectin expression in our study promoting 3D cell migration and invasion after depletion of PRP4K in MCF10A cells (Fig 2 and 3).Finally, in a previous study we observed the upregulation of Zeb1 following depletion of PRP4K in the epithelial-like breast cancer cell line MCF7 that also occurred without loss of E-cadherin (21). Although more variable in the current study and most consistent in MDA-MB-231 cells, we also observed increased expression of Zeb1 in our cell lines after depletion of PRP4K with at least one shRNA (Supplemental Figure S2). We also observed that protein expression changes were not well correlated with mRNA expression in PRP4K depleted cells (Supplemental Figure S3). This may be expected given the well-established molecular heterogeneity of EMT factor expression in cancer, where a variety of mesenchymal and epithelial gene expression patterns are observed (40). This heterogeneity is also a reflection of post-transcriptional control of protein expression during EMT that likely is responsible for differences in EMT factor mRNA and protein expression (41), which we also observe for PRP4K protein regulation during EMT induced by eIF3e depletion (discussed below). Overall, these data are consistent with depletion of PRP4K inducing a partial EMT state in normal mammary epithelial cell lines (Figure 11A).

**Figure 4.**
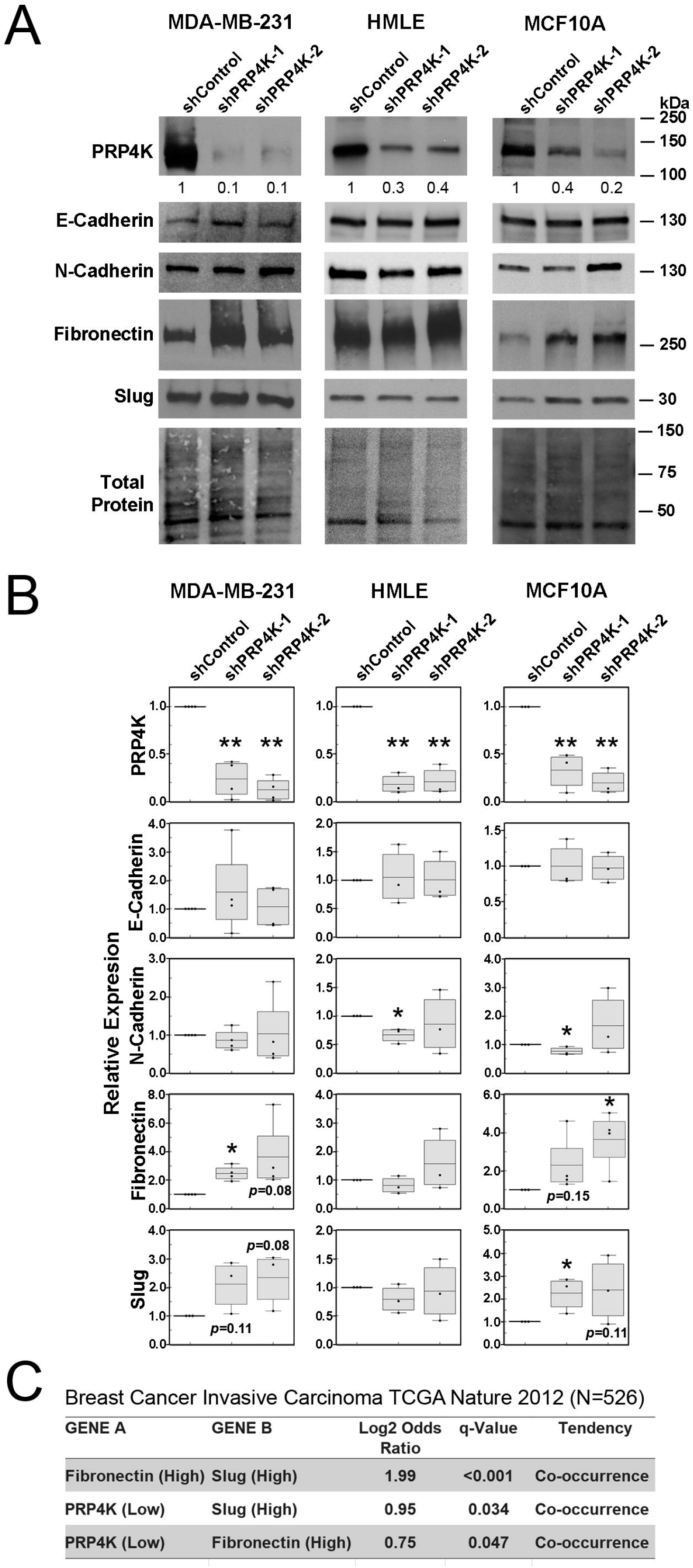
Loss of PRP4K induces changes in mesenchymal markers without changes in E-cadherin. **A)** Whole-cell lysates of MDA-MB-231, HMLE, and MCF10A control (shCtrl) and PRP4K depleted (shPRP4K-1 and shPRP4K-2) cells were prepared and subject to Western blot analysis of epithelial (E-cadherin,) and mesenchymal (Slug, N-cadherin, Fibronectin) proteins as indicated. Ratios of protein level, normalized to total protein are shown below each row of Western blots relative to either shCtrl levels (set to 1) or lowest expression in PRP4K-depleted cells. **B**) Quantification of protein expression in panel (A) using densitometry of protein bands relative to actin or total protein. Relative expression values to control (set to 1) are represented as a box and whisker plot, with the mean and upper and lower quartiles indicated by the bounding box. N = 3 or 4 replicates (as indicated by data points) and error bars = SD. Significance was determined by a t-test (two-tailed). *p<0.05. **C**) Analysis of TCGA Breast Invasive Carcinoma mRNA expression data (31). The Logrank test was used to determine the co-occurrence of low PRP4K (*PRPF4B*)(i.e. < −1 SD relative to diploid samples) with high expression of fibronectin (*FN1*) or Slug (*SNAI2*)(i.e. > 1 SD relative to diploid samples). Log2 Odds Ratio is shown, and p-value estimates of significance were adjusted for false discovery rate and reported as q-values. N =526 tumours.

Since the induction of EMT can trigger gene expression changes that promote aggressive cancer phenotypes (1, 4), we next examined whether the induction of EMT affected the expression of PRP4K. Employing two different methods of EMT-induction involving either the exposure of cells to media containing WNT-5a and TGFβ1 (EMT-induction media (33), Figure 5), or depletion of eIF3e (27) (Figure 7), we determined that PRP4K protein expression is negatively regulated by the induction of complete EMT (as marked by dramatic loss of E-cadherin and upregulation of N-cadherin, Figure 5B and 7B) in MCF10A and HMLE cells. We found that EMT-induction media caused a reduction in PRP4K mRNA expression in MCF10A cells and did not affect PRP4K protein translation (Figure 6); however, this was not true for EMT induced by eIF3e depletion. In contrast, eIF3e depletion had no effect on PRP4K mRNA levels (Figure 7C), transcript turn-over (Figure 8A) or protein stability (Figure 8B) but rather reduced PRP4K protein translation by ~40% (Fig 8C). This was an intriguing result that leads us to hypothesize that PRP4K may be negatively regulated by different mechanisms depending on the EMT-inducing stimulus; with TGF-β1-induced EMT negatively regulating PRP4K at the transcriptional level (Figure 11A), whereas EMT triggered by eIF3e depletion results in inhibition of PRP4K translation (Figure 11B). One limitation in generalizing changes in PRP4K translation relative to EMT is that we cannot exclude the possibility that depletion of eIF3e alone may be responsible for reduced translation of PRP4K, rather than EMT *per se*.

**Figure 5.**
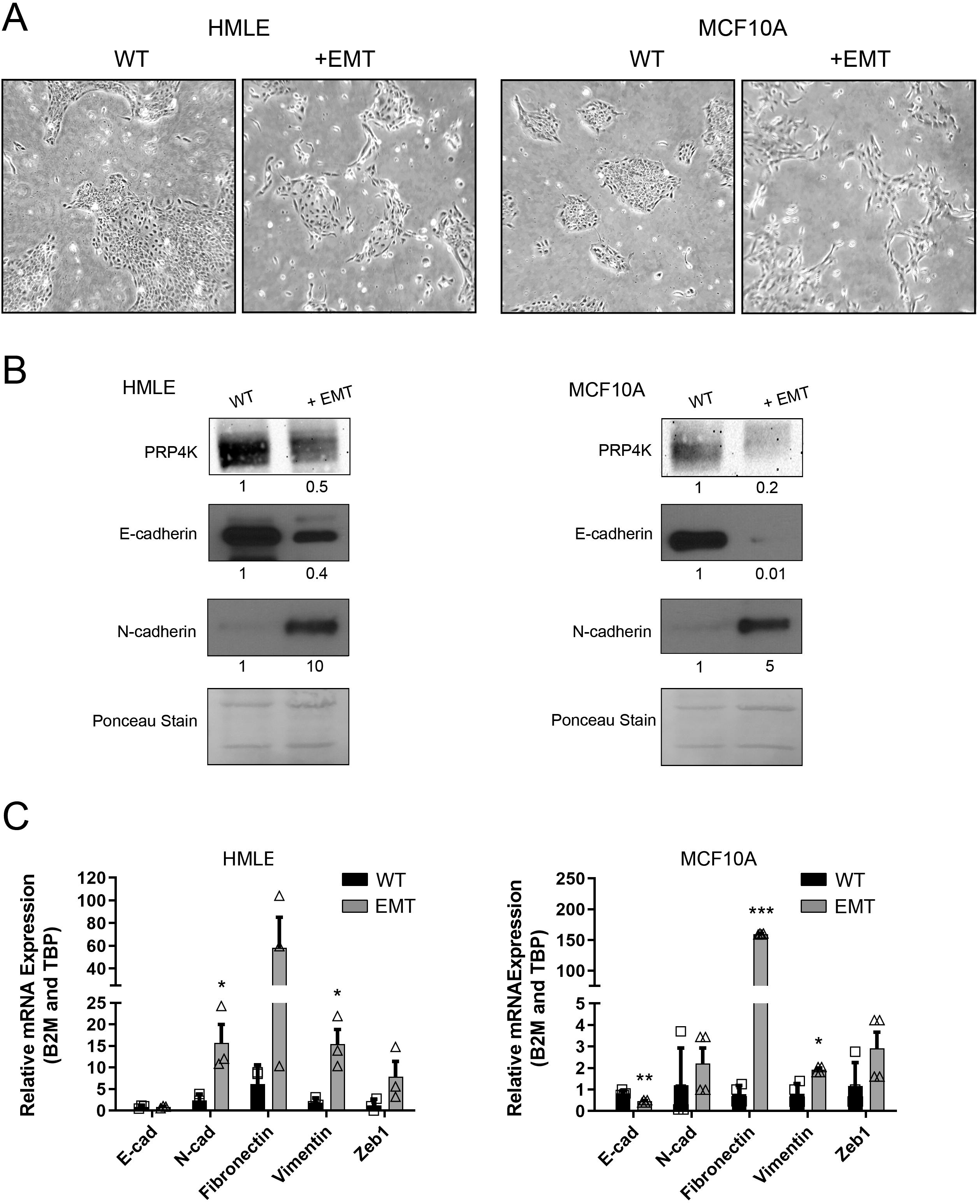
The induction of EMT decreases PRP4K protein expression in HMLE and MCF10A cells. **A)** Phase contrast micrographs of MCF10A and HMLE cells treated with (+EMT) or without (WT) EMT-induction media for 5 days. **B**) Western blot analysis of PRP4K protein expression from whole cell lysates prepared from cells treated with EMT-induction media as in A. **C**) Quantitative PCR of cDNA prepared from cells treated with EMT-induction media as in A, with data normalized to at least two reference genes as indicated. N = 3, error bars= SEM. Significance was determined by a t-test. *=p<0.05.

**Figure 6.**
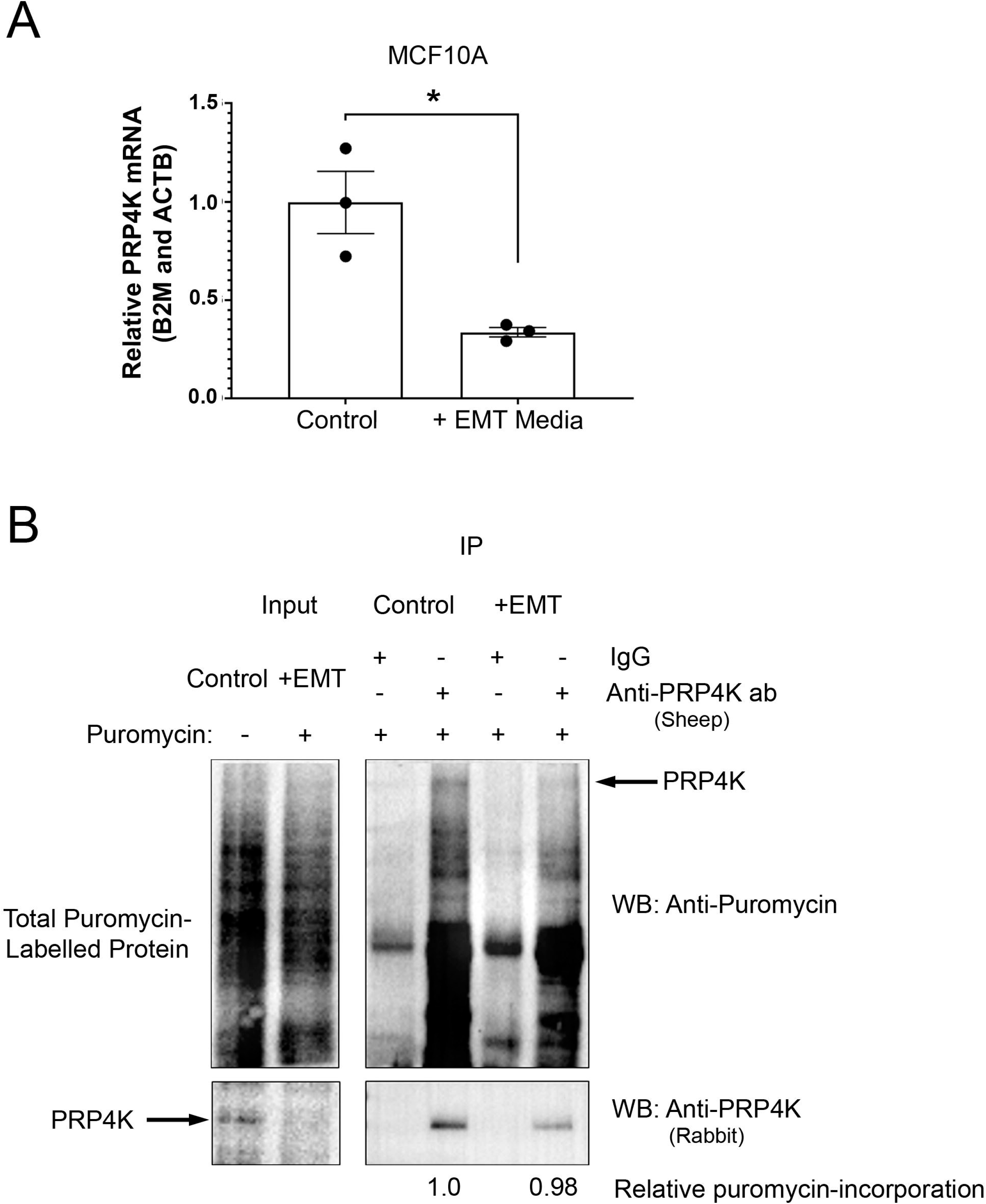
The induction of EMT by EMT-induction media negatively regulates PRP4K mRNA expression in MCF10A cells. **A)** Quantitative PCR analysis of PRP4K gene (*PRPF4B*) mRNA expression in untreated MCF10A (Control) cells or those treated with EMT-induction media (+ EMT) for 5 days, with data normalized to at least two reference genes as indicated. N = 3, error bars= SEM. Significance was determined by a t-test (two-tailed). *=p<0.05. **B**) Puromycin incorporation assay of untreated MCF10A (Control) and EMT-induction media treated cells as in panel (A). After EMT-induction, cells were grown for 24 h without puromycin selection and then pulsed with puromycin prior to Western blot analysis of total cell lysates (left) or following immunoprecipitation (IP) of PRP4K and detection of puromycylation of proteins and PRP4K (right). Relative puromycin incorporation normalized by total PRP4K in the IP is shown.

Partial EMT states can promote cancer cell survival, increased plasticity and collective cell migration and invasion by cancer cells that increases their metastatic potential and could contribute to worse clinical outcomes (4, 5). Metastasis in ovarian cancer is associated with collective cell migration and invasion (42), and previously we demonstrated that high-grade serous ovarian cancer (HGSC) patients harboring tumours with low PRP4K expression had significantly worse clinical outcomes; with low PRP4K protein expression correlating with both acquisition of taxane chemotherapy resistance as well as worse overall survival (21). Conversely, amplification of the PRP4K gene *PRPF4B* on human chromosome 6 is associated with significantly better overall survival in ovarian cancer (21), data that together strongly indicates that PRP4K is a tumour suppressor in HGSC. In breast cancer, we have documented taxane resistance in both ER-positive and TNBC breast cancer cell lines after PRP4K depletion (23), and Cho and colleagues have reported a significant correlation between low PRP4K expression in breast tumours and worse overall survival in patients with TNBC (26). However, in a second study by Koedoot *et al*.,(38) high expression of PRP4K correlated with worse overall survival in patients with TNBC. One caveat in interpreting these contradictory results is that different cohorts of breast cancer patients were used in the two studies. TNBC is a rather broad and heterogeneous subtype of breast cancer (43), and thus future studies of specific molecular subtypes of TNBC (including aberrant signaling pathways) may be required to better define the clinical impact of low PRP4K expression.

One important signaling pathway for cancer development and progression is the Hippo-Yap axis (44). The Hippo pathway is known to negatively regulate the transcription factors YAP and TAZ, which are upregulated in many cancer types and control transcriptional programs that promote EMT and increased metastatic potential (45–47). Cho and colleagues identified YAP as a substrate of PRP4K, which negatively regulates YAP by promoting its phosphorylationdependent shuttling from the nucleus to the cytoplasm (26). We observed that the induction of EMT by eIF3e depletion resulted in reduced expression of PRP4K (Figure 7B), which correlated with increased nuclear versus cytoplasmic localization of YAP (Figure 9) as well as increased expression of YAP target genes in MCF10A mammary epithelial cells (Figure 10A)(Summarized in Figure 11B). These data are reminiscent of the activation and nuclear accumulation of YAP/TAZ previously seen in the context of TGF-β1-mediated EMT in mouse mammary epithelial cell line NmuMG (48), which suggests that nuclear accumulation of YAP/TAZ is likely a general feature of EMT. We also observed that overexpression of PRP4K, to complement PRP4K loss in eIF3e-depleted cells, shifts the localization of YAP toward accumulation in the cytoplasm. This cytological change in YAP localization correlated with inhibition of YAP-target gene expression (*ANKRD1, CTGF, CYR61*) in cells with reduced eIF3e that have overexpression of exogenous PRP4K compared to eIF3e-depleted MCF10A cells transfected with a non-expressing plasmid (i.e. pBluescript; Figure 10). These results are consistent with the findings of Cho *et al*., (26) that PRP4K can regulate the nucleo-cytoplasmic shuttling of YAP and YAP-target gene expression, and as such, our data support a role for PRP4K loss in promoting increased YAP activity and nuclear accumulation during EMT.

**Figure 7.**
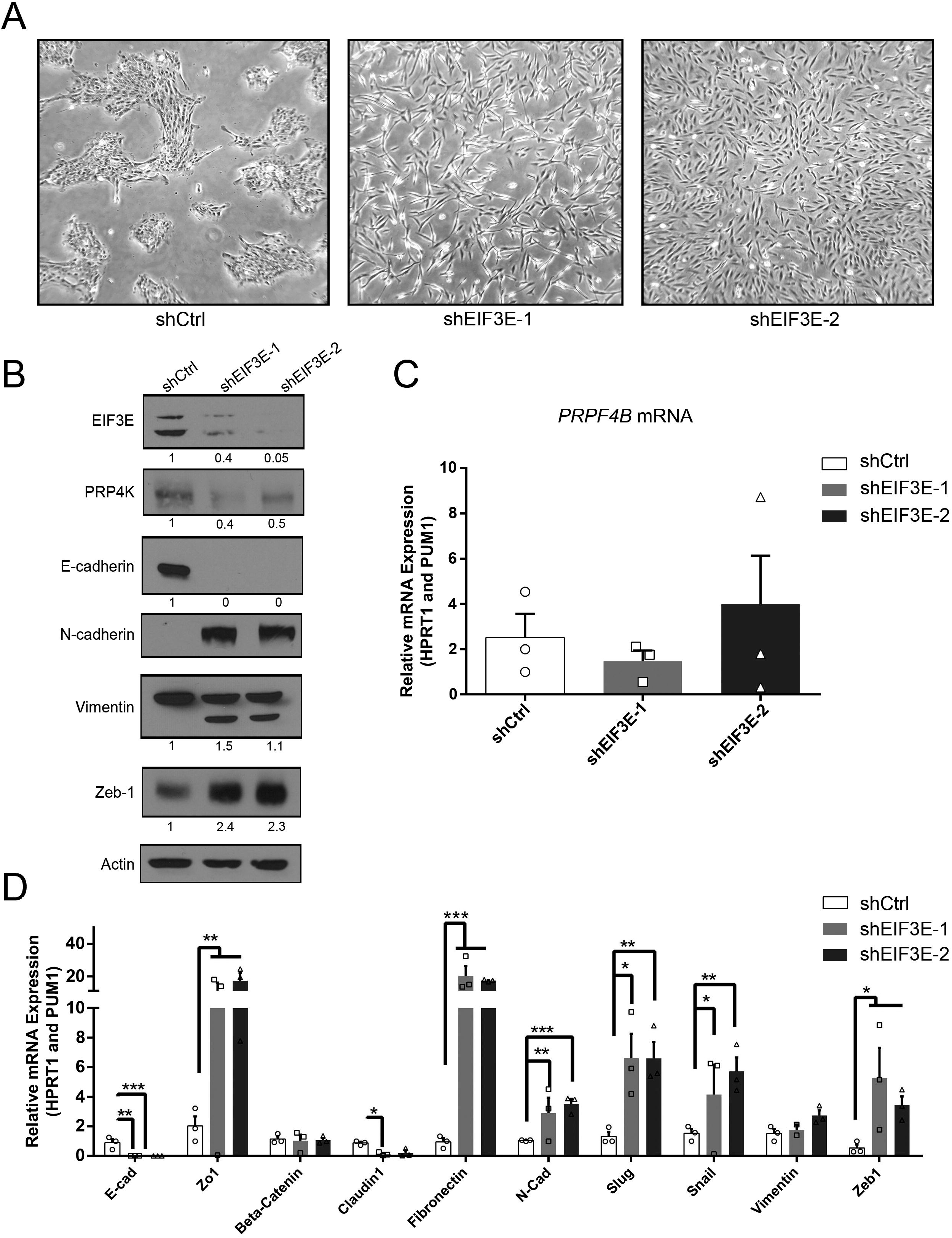
The induction of EMT by depletion of eIF3e decreases PRP4K protein expression in MCF10A cells. **A)** Phase contrast micrographs of control (shCtrl) and eIF3e-depleted cells (shEIF3E-1 and shEIF3-2). **B**) Western blot analysis of whole cell lysates of MCF10A shCtrl and shEIF3E cells for markers of EMT as indicated. **C**) Quantitative PCR of PRP4K gene (*PRPF4B*) mRNA expression in MCF10A shCtrl and shEIF3E cells. **D**) Quantitative PCR analysis of EMT-associated gene expression (as indicated) in MCF10A shCtrl and shEIF3E cells. N = 3, error bars= SEM. Significance was determined by a t-test. *=p<0.05, **=p<0.01, ***=p<0.001.

**Figure 8.**
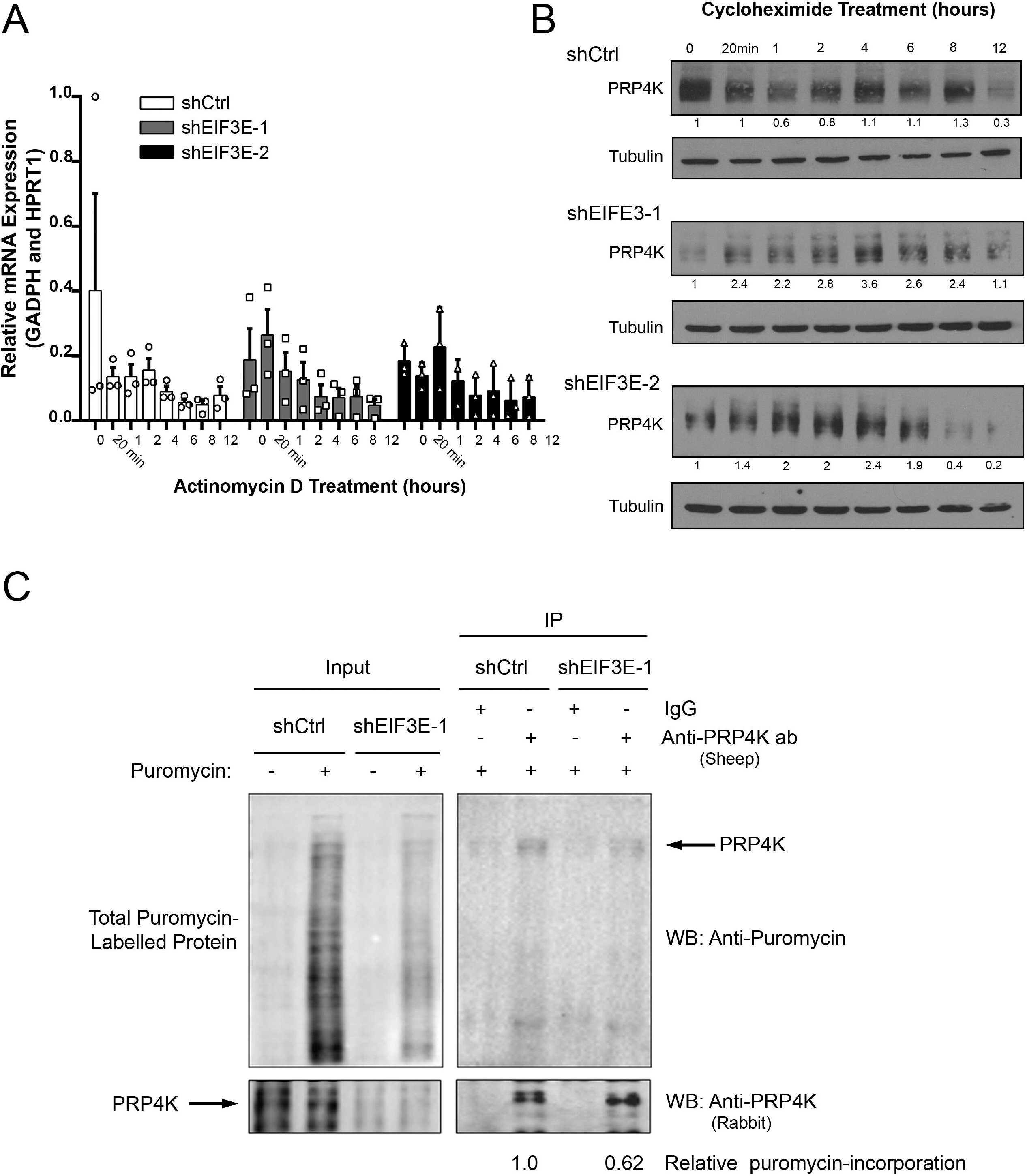
The induction of EMT by depletion of eIF3e negatively regulates the translation of PRP4K in MCF10A cells. **A)** Quantitative PCR of PRP4K gene (*PRPF4B*) mRNA expression in control (shCtrl) and eIF3e-depleted (shEIF3E-1 and shEIF3-2) MCF10A cells treated with 10 μg/mL Actinomycin D for the indicated time periods, with data normalized to at least two reference genes as indicated. N = 3, error bars= SEM. Significance was determined by a one-way ANOVA. **B**) Western blot analysis of whole cell lysates of MCF10A shCtrl and shEIF3E cells treated with 50μg/mL cycloheximide for the indicated time periods. **C**) Puromycin incorporation assay of MCF10A shCtrl and shEIF3E cells. Cells were grown for 24 h without puromycin selection and then pulsed with puromycin prior to Western blot analysis of total cell lysates (left) or following immunoprecipitation (IP) of PRP4K and detection of puromycylation of proteins and PRP4K (right). Relative puromycin incorporation normalized by total PRP4K in the IP is shown.

**Figure 9.**
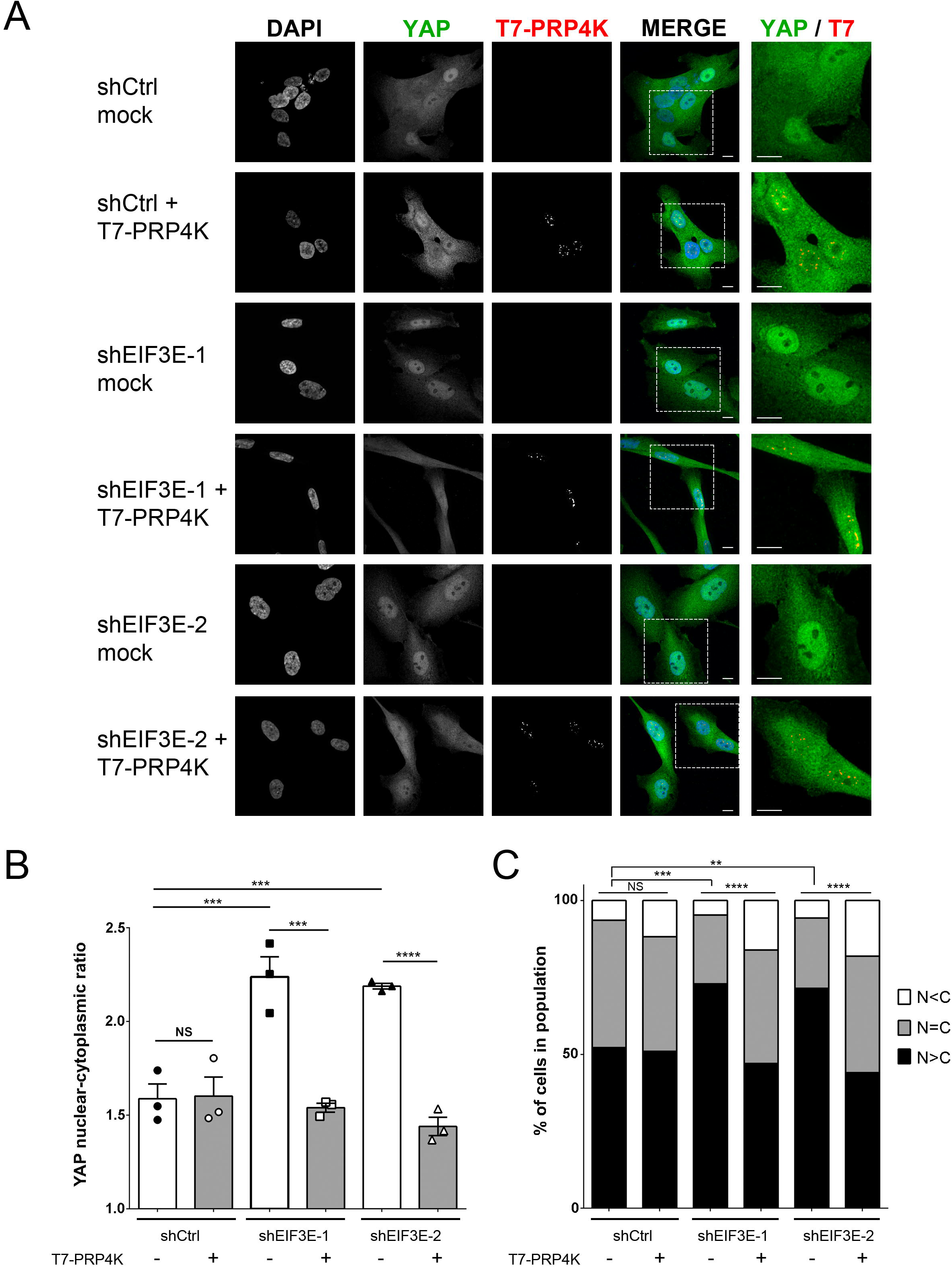
Depletion of eIF3e in MCF10A cells increases YAP nuclear localization and is reversed by overexpression of PRP4K. **A)** Immunofluorescence images of control (shCtrl) and eIF3e-depleted MCF10A cells (shEIF3E-1 and shEIF3-2) transfected with pBluescript SK-(mock) or T7-tagged PRP4K (T7-PRP4K) plasmids. An enlarged region of each merged micrograph, bound by the white box, is shown in the far right panel. DAPI=blue, YAP=green, T7=red. Scale bars = 10 microns. **B**) Histogram of YAP nuclear-cytoplasmic ratios for shCtrl and shEIF3E cells transfected with pBluescript SK-(-) or with (+) T7-tagged PRP4K. N = 3, error bars=SEM. Significance was determined by a one-way ANOVA. **C**) Graph showing the percentage of cells in B with N<C, N=C and N>C expression of YAP. Significance was determined by a Fisher’s exact test. **=p<0.01, ***=p<0.001, ****=p<0.0001.

**Figure 10.**
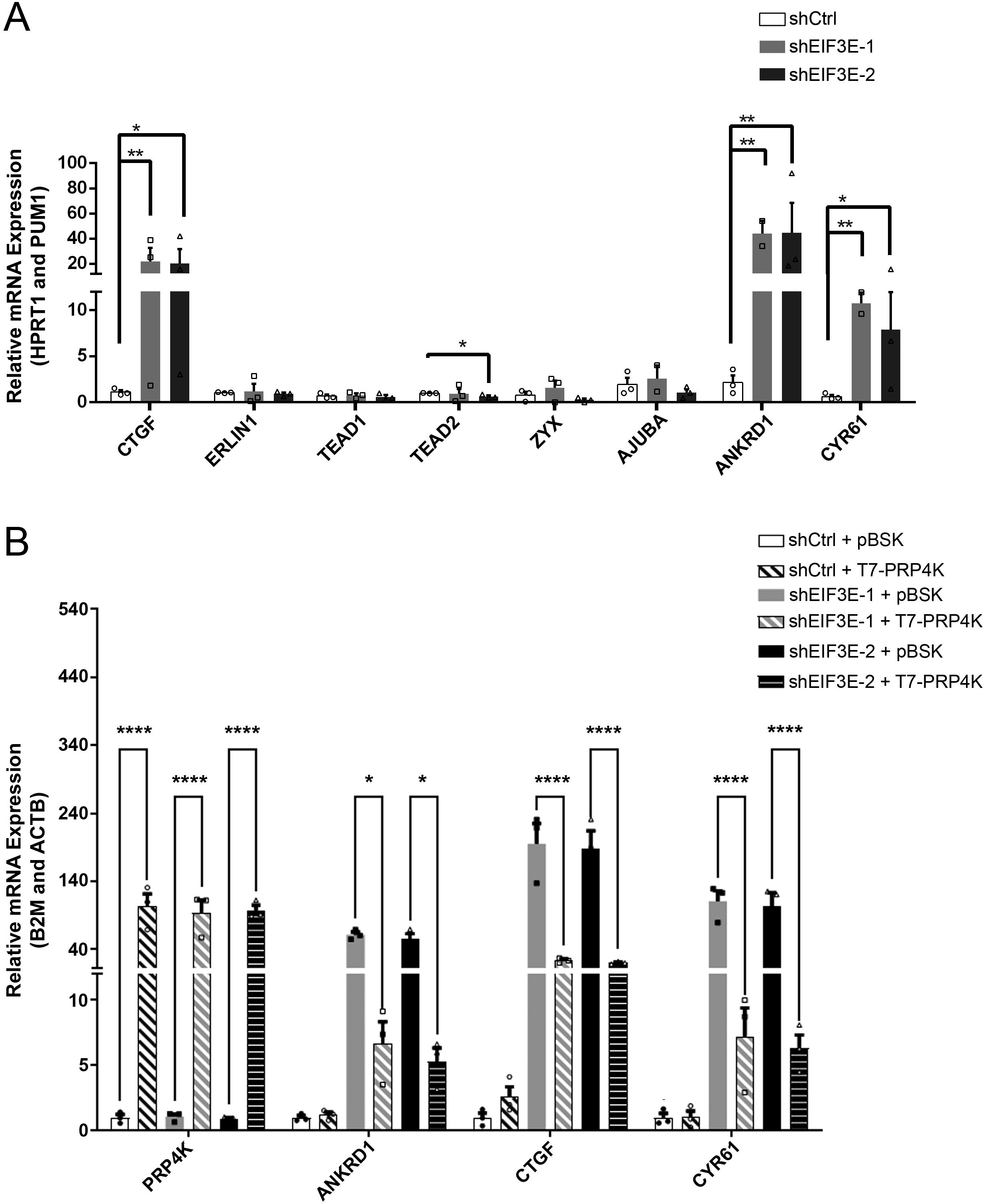
Overexpression of PRP4K decreases YAP target genes in eIF3e-depleted MCF10A cells. **A)** Quantitative PCR analysis of cDNA prepared from control (shCtrl) and eIF3e-depleted cells (shEIF3E-1 and shEIF3-2), with data normalized to at least two reference genes as indicated. N = 3, error bars= SEM. Significance was determined by a one-way ANOVA. **B**) Quantitative PCR analysis of cDNA prepared from control (shCtrl) and eIF3e-depleted cells (shEIF3E-1 and shEIF3-2) transfected with pBluescript SK-(pBSK) or T7-tagged PRP4K, with data normalized to at least two reference genes as indicated. N = 3, error bars= SEM. Significance was determined by a one-way ANOVA. *=p<0.05, **=p<0.01, ****=p<0.0001.

**Figure 11.**
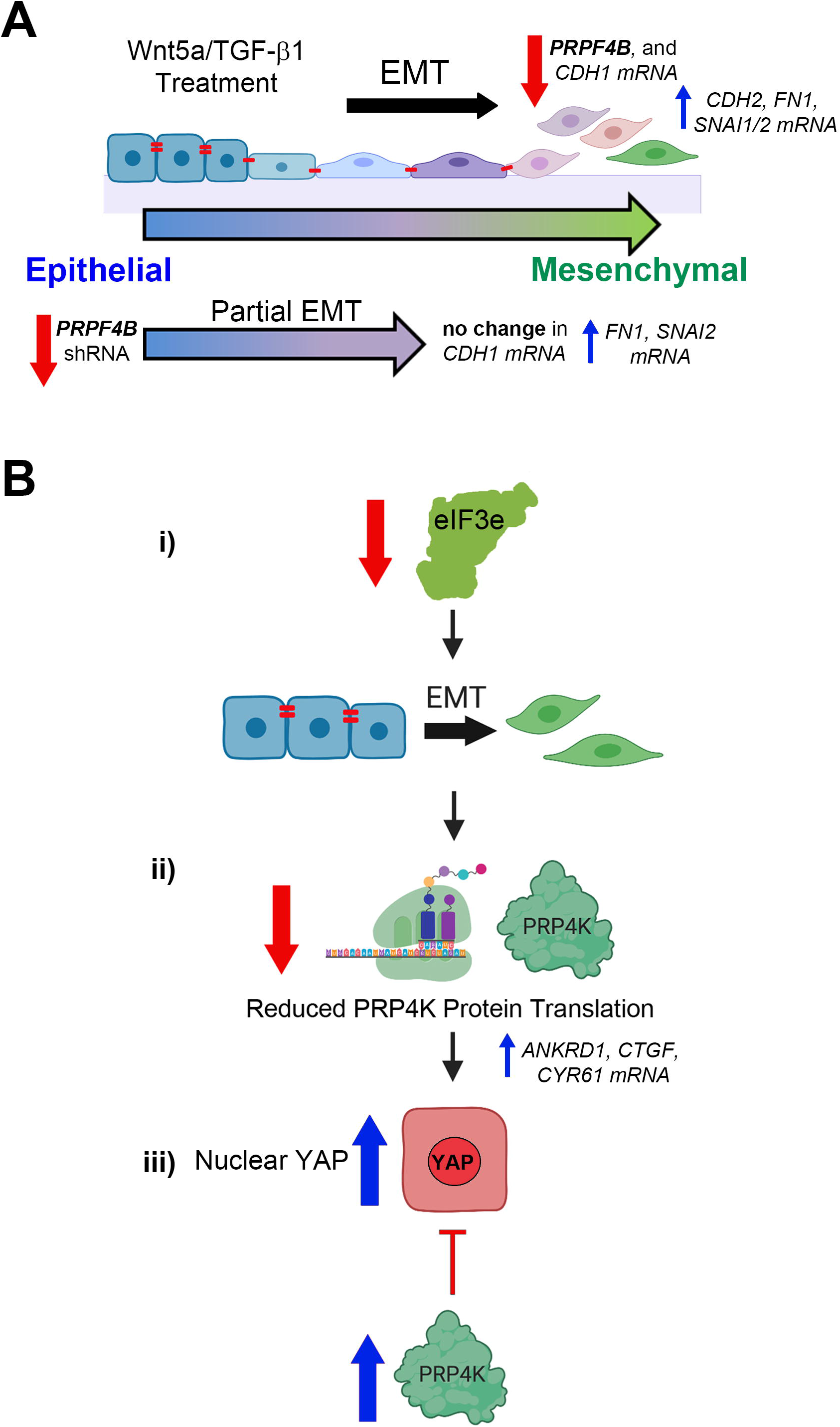
Overview of the key findings on the relationship between PRP4K and EMT. **A)** Epithelial-to-mesenchymal transition (EMT) occurs over a spectrum of phenotypes from full to partial EMT. EMT-induction media containing WNT-5a and TGF-β1 can induce full EMT and a marked reduction in PRP4K gene (PRPF4B) mRNA expression, which is correlated with reduced E-cadherin (*CDH1*) and increased mesenchymal gene expression including N-cadherin (*CDH2*), fibronectin (FN1), Snail (*SNAI1*) and Slug (*SNAI2*) However, depletion of PRP4K by shRNA in the mammary epithelial cells promotes a partial EMT phenotype, marked by no change in E-cadherin, increased mesenchymal gene expression (*FN1, SNAI2*) and increases cell invasion potential. **B**) Model of the effects of eIF3e depletion on PRP4K levels and YAP localization in MCF10A cells. i) Depletion of eIF3e induces EMT. ii) EMT induction through eIF3e depletion reduces PRP4K protein translation, which is correlated with increased YAP activity and nuclear localization. iii) Overexpression of PRP4K inhibits the nuclear localization of YAP and expression of YAP-target genes. Elements of this figure were created with BioRender.com.

In summary, this study provides further support for PRP4K as a haploinsufficient tumour suppressor, which we demonstrate here to be negatively regulated by EMT, and that when depleted can induce partial EMT to promote more aggressive cancer phenotypes including increased cell invasion.

## ABBREVIATIONS

EMT: Epithelial-to-mesenchymal transition
MET: mesenchymal-to-epithelial transition
PRP4K: pre-mrna processing factor 4 kinase
shRNA: small short hairpin RNA
TAZ: Tafazzin
2D: Two dimensional
3D: three dimensional
TGF-β1: Transforming growth factor-beta 1
TNBC: triple negative breast cancer
WNT5a: Wnt oncogene analog 5a
YAP: Yes-associated protein

## ACKNOWLEDGMENTS

This work was funded by a Breast Cancer Society/QEII Foundation operating grant awarded to GD and an Operating Grant (#24135) from the Cancer Research Society and the Beatrice Hunter Cancer Research Institute (BHCRI) awarded to SML. GD and SML are Senior Scientists of the BHCRI, and both AC and LEC were supported by Cancer Research Training Program awards from the BHCRI with funds provided by CBCF – Atlantic, The Canadian Cancer Society, Nova Scotia Division as part of The Terry Fox Foundation Strategic Health Research Training (STIHR) Program in Cancer Research at the Canadian Institutes of Health Research (CIHR). SM is supported by a CGSM studentship from Natural Sciences and Engineering Research Council of Canada (NSERC), a Predoctoral Fellowship from the Killam Trusts, and a Scotia Scholar award from the Nova Scotia Health Research Foundation.

## AUTHOR CONTRIBUTIONS

G. Dellaire and S.M. Lewis obtained funding and designed the research; L.E. Clarke, A. Cook, S. Mathavarajah, E. Habib, J. Salsman, C. Van Iderstine, A. Bera, and M. Bydoun performed research and analyzed data; L.E. Clarke and G. Dellaire are responsible for data interpretation and wrote the manuscript; and all authors contributed to editing the final manuscript.

