## Supplemental Figures for "Haploinsufficient tumour suppressor PRP4K is negatively regulated during epithelial-to-mesenchymal transition"

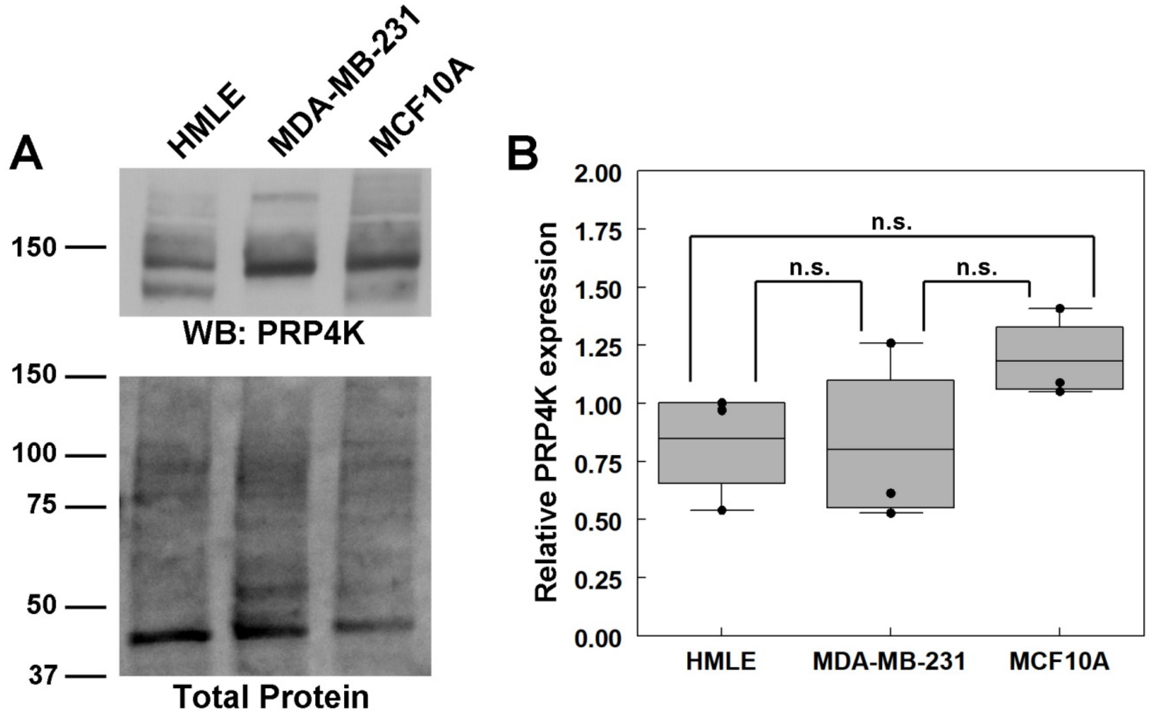


**Supplemental Figure S1. Relative PRP4K protein expression in MDA-MB-231, HMLE and MCF10A cells. A**) Whole-cell lysates of MDA-MB-231, HMLE, and MCF10A cells were prepared and subject to Western blot analysis of PRP4K protein levels. Ratios of protein level, normalized to total protein are shown below each row of Western blots relative to either shCtrl levels (set to 1). **B**) Quantification of PRP4K protein expression in panel (A) using densitometry of protein bands relative to total protein. Relative expression values are shown relative to the highest expression of PRP4K in HMLE cells (set to 1) and depicted as a box and whisker plot, with the mean and upper and lower quartiles indicated by the bounding box. N = 3 or 4 replicates (as indicated by data points) and error bars = SD. N=3 and error bars = SD. N.S. = not significant.

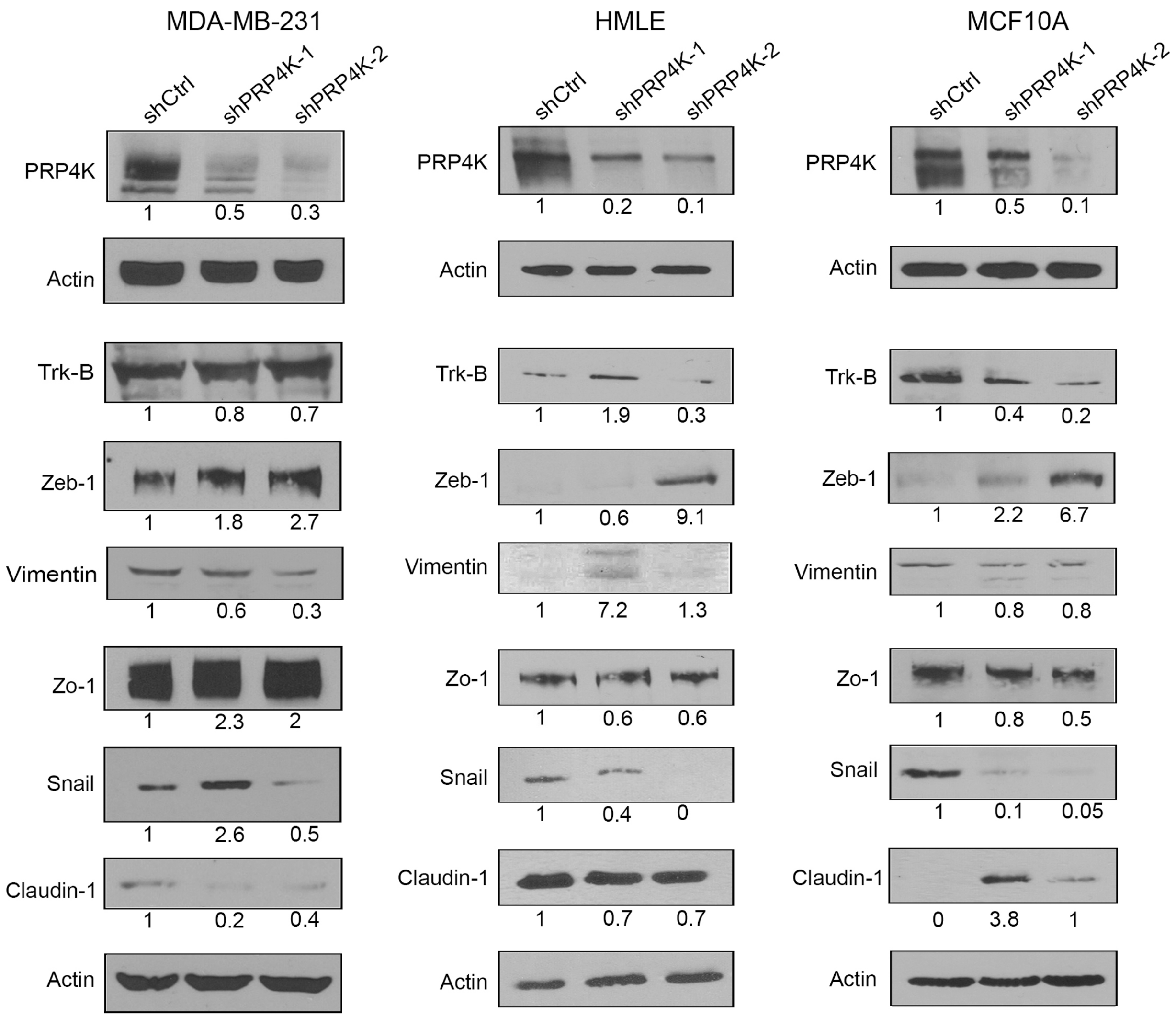


**Supplemental Figure S2. Changes in epithelial and mesenchymal protein expression associated with PRP4K depletion in MDA-MB-231, HMLE and MCF10A cells.** Whole-cell lysates of MDA-MB-231, HMLE, and MCF10A control (shCtrl) and PRP4K depleted (shPRP4K-1 and shPRP4K-2) cells were prepared and subject to Western blot analysis of epithelial (Zo-1, claudin-1) and mesenchymal (Snail, Zeb1, Trk-B, N-cadherin, Vimentin, Fibronectin) proteins as indicated. Ratios of protein level, normalized to actin, are shown below each row of Western blots relative to either shCtrl levels (set to 1) or lowest expression in PRP4K-depleted cells.

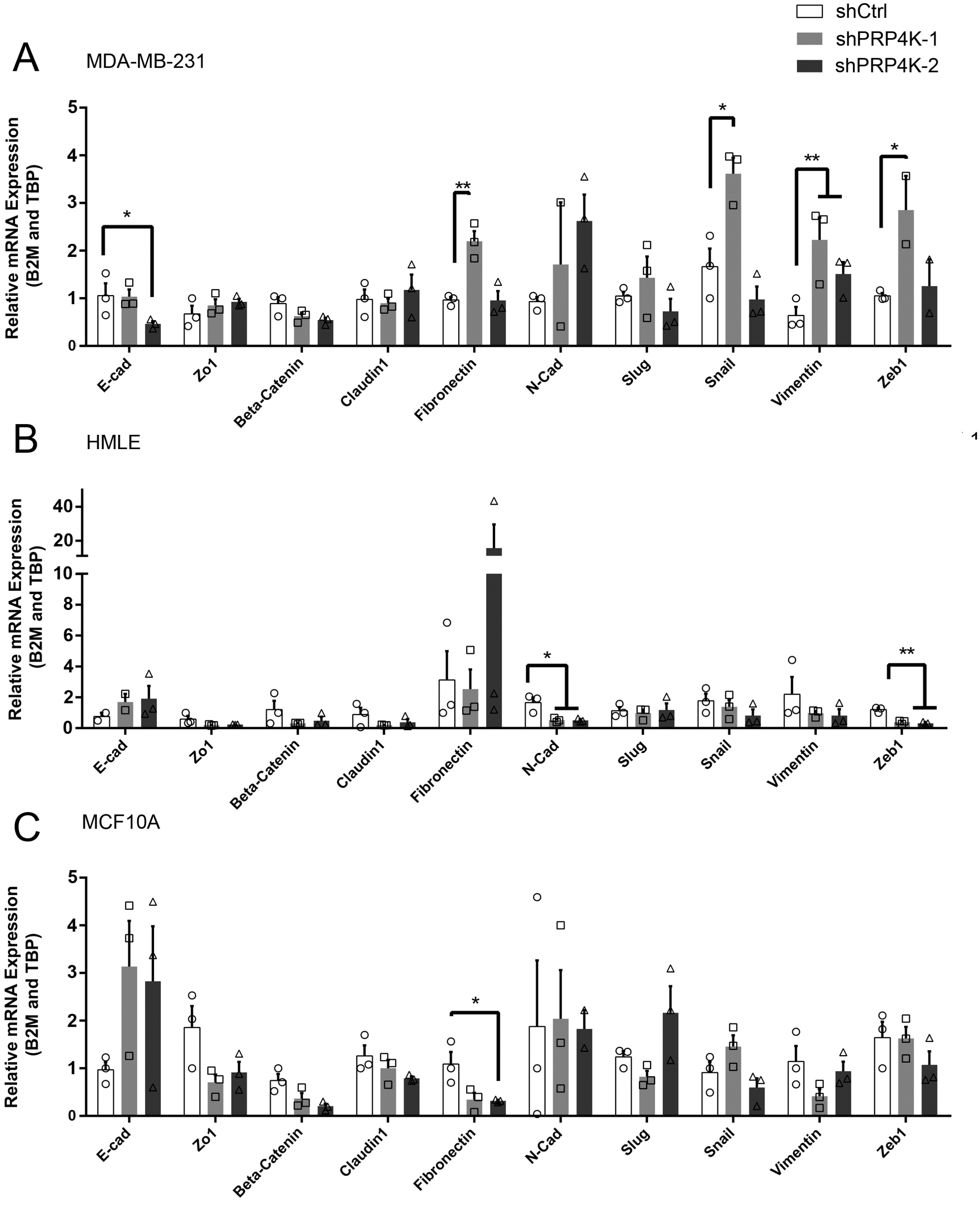


**Supplemental Figure S3. Changes in epithelial and mesenchymal gene expression associated with PRP4K depletion in MDA-MB-231, HMLE and MCF10A cells. A-C**) RNA was isolated from the shCtrl and shPRP4K cells (cell lines indicated) and then reverse-transcribed to cDNA. Reverse transcription quantitative PCR (RT-qPCR) was performed on the cDNA samples and all data was normalized against two reference genes (*B2M* and *TBP*). N = 3, error bars= S.E.M. Significance was determined by a t-test. *p<0.05, **p<0.01
